# Evolution of the CRISPR-Cas9 defence system in *Mycoplasma gallisepticum* following colonization of a novel bird host

**DOI:** 10.1101/2023.03.14.532377

**Authors:** Thomas Ipoutcha, Iason Tsarmpopoulos, Géraldine Gourgues, Vincent Baby, Paul Dubos, Geoffrey E. Hill, Yonathan Arfi, Carole Lartigue, Patricia Thebault, Camille Bonneaud, Pascal Sirand-Pugnet

## Abstract

CRISPR-Cas systems are bacterial defences that target bacteriophages and mobile genetic elements. How these defences evolve in novel host environments remains, however, unknown. We studied the evolution of the CRISPR-Cas system in *Mycoplasma gallisepticum*, a bacterial pathogen of poultry that jumped into a passerine host ∼30 years ago. Over the decade following the host shift, all isolates displayed a functional CRISPR-Cas system were found not only to harbour completely new sets of spacers, but the DNA protospacer adjacent motif (PAM) recognised by the main effector MgCas9 was also different. These changes in CRISPR-Cas diversity and specificity are consistent with a change in the community of phages and mobile elements infecting *M. gallisepticum* as it colonised the novel host. In the years following the host shift, we also detected a gradual rise in isolates displaying non-functional MgCas9. After 12 years, all circulating isolates harboured inactive forms only. This loss of CRISPR-Cas function comes at a time when the passerine host is known to have evolved widespread resistance, which in turn drove the evolution of increasing *M. gallisepticum* virulence through antagonistic coevolution. Such striking concordance in the rise of inactivated forms of CRISPR-Cas and the evolution of host resistance suggests that inactivation of the CRISPR-Cas system was necessary for enabling adaptive bacterial responses to host-driven selection. We highlight the need to consider both host and pathogen selection pressures on bacteria for understanding the evolution of CRISPR-Cas systems and the key factors driving the emergence of a pathogenic bacterium in a novel host.

**Data summary:** The authors confirm all supporting data and protocols have been provided within the article or through supplementary data files available in the online version of this article. GenBank accession numbers of all publicly available *M. gallisepticum* genomes are listed in Table S3. Sequences of the CRISPR locus of other strains are also provided in Table S3.

**Impact statement:** Mycoplasma are minimal bacteria involved in many diseases affecting humans and a wide diversity of animals. In this paper, we report the evolution of the Type II CRISPR-Cas system of the bird pathogen, *Mycoplasma gallisepticum*, following an host jump from its original poultry host into its novel house finch host in the early 90’s. Instances in which bacterial pathogens have been documented to jump into and subsequently adapt to a new host are rare, and the well documented case of *M. gallisepticum* is a unique model to evaluate the effect of any dramatic host environmental change on bacterial CRISPR-Cas defence systems. First, we performed in silico analyses on an extended set of 98 *M. gallisepticum* genomes to better understand the evolution of the CRISPR-Cas9 system in the novel finch host. We documented several evolutionary events leading to the drastic divergence of spacer sets present in poultry and house finch arrays, as well as the progressive inactivation of the CRISPR-Cas system after 12 years in the novel finch host. Second, using in vitro and in vivo assays, we demonstrated that the evolution of the MgCas9 PI domain, involved in the protospacer adjacent motif (PAM) recognition has led to a major change in the defence system, with a modification of the recognized PAM in the novel host. Such radical change in the CRISPR-Cas defence system of *M. gallisepticum* may have implications for the its rapid adaptation to its novel host. Together, our results highlight the need to consider not only the host-driven selection pressures a bacterium experiences, but also the complex interplay between phages and defence systems for better understanding the key factors driving the emergence of a pathogenic bacterium in a novel host.

## Introduction

Clustered Regularly Interspaced Short Palindromic Repeats (CRISPR) systems are defences found in most prokaryotes that effectively protect cells against the constant threat of phages and other sources of invading DNAs. The flexibility and effectiveness of this defence system relies on the continuous acquisition of new spacers to the CRISPR array (Barrangou et al., 2007), which entails substantial maintenance costs. As a consequence, inactivation and disappearance of CRISPR systems from prokaryote genomes is predicted to occur when benefits of this defence system are relaxed (Koonin, 2019; Makarova et al., 2020). Ultimately, gain or loss events of active CRISPR systems will be shaped both by cost/benefit ratios (Dimitriu et al., 2020) and by the complex ecological pressures experienced by prokaryotes. For pathogenic bacteria, the strongest ecological pressures arise from interactions with hosts. Changes in host-driven selection pressures may occur, for instance, as hosts evolve in response to pathogens, but should be particularly dramatic when a prokaryotic parasite colonizes a novel host. While invasion of a novel host is known to broadly impact bacterial genomes, how such host shifts shape bacterial defence systems against phages and other parasitic DNAs remains largely unstudied.

Instances in which bacterial pathogens have been documented to jump into and subsequently adapt to a new host species are rare (Bonneaud et al. 2019), limiting opportunities to test the effect of host environmental change on bacterial CRISPR systems. One model in which the adaptation of a bacterium to a novel host can be studied is the *Mycoplasma gallisepticum* / house finch (*Haemorhous mexicanus*) system. *M. gallisepticum* is a bacterial pathogen that is a common in domestic chickens (*Gallus gallus*), a gallinaceous bird. It jumped into a very distantly related and widely distributed North American passerine, the house finch. This novel host shift was documented in 1994 by wildlife disease experts (Ley et al., 1996) and a mid-1990s host-shift date has been confirmed by genome analyses of *M. gallisepticum* isolates collected from chickens and finches. *M. gallisepticum* lineages subsequently collected across the range of the house finches indicate that all *M. gallisepticum* circulating in house finch populations are derived from this single host-shift event (Delaney et al., 2012; Tulman et al., 2012).

*M. gallisepticum* is an avian pathogen that belongs to the class *Mollicutes* (May et al., 2014), a group of minimal bacteria characterized by a rapid evolution through drastic genome reduction from a Gram-positive ancestor. Such genetic losses are partially counter-balanced by horizontal gene transfers (HGT) and exposure to Mobile Genetic Elements (MGEs) acting as potential sources of genetic diversity and genome reorganization (Citti et al., 2018; Sirand-Pugnet et al., 2007). Despite an evolutionary tendency toward genome minimization, most mycoplasmas have retained various defence systems against phages and invading DNAs. Indeed, mycoplasmas are the second group of bacteria with the greatest number of Restriction-Modification systems (Oliveira et al., 2014), and are also the free-living bacteria with the smallest genome in which CRISPR-Cas systems have been described (Ipoutcha et al., 2019).

Changes in the CRISPR locus of *M. gallisepticum* have previously been reported following the host shift into house finches (Delaney et al., 2012; Dhondt et al., 1998). Indeed, a comparison of *M. gallisepticum* isolates obtained from ancestral poultry and novel house finch hosts showed a complete change in the spacer set, with no elements being shared between poultry and house finch strains (Delaney et al., 2012). House finch isolates also displayed a loss of spacers over time, as well as a degradation of their CRISPR locus. While these patterns suggest both a slowing down in the recruitment of new spacers and even an evolution towards this defence system becoming non-functional, further work is required to test whether these evolutionary changes have persisted over time in the new passerine host. Making sense of observed changes in the house finch CRISPR also requires functionally characterising other changes that have occurred in the CRISPR-Cas system, and particularly in *M. gallisepticum* Cas9 (MgCas9). Previous research has shown that Cas9 endonuclease is the main effector of type-II CRISPR systems, being involved in both the acquisition of new spacers and the double-strand DNA cleavage of target DNAs (Heler et al., 2015; Wei et al., 2015). In both processes, Cas9 selects functional spacers by recognizing their protospacer adjacent motif (PAM) sequence immediately downstream. The change of spacer set that has accompanied *M. gallisepticum* host jump therefore calls into question the evolution of MgCas9 PAM specificity as a possible driver.

To better understand the changes in the *M. gallisepticum* CRISPR system that took place during the colonization of the novel avian host, we conducted a functional characterisation using a two-pronged approach. First, we performed *in silico* analyses on an extended set of *M. gallisepticum* isolates (79 collected from 1994 to 2015) to better understand evolutionary changes in CRISPR-Cas9. We identified several events leading to the remarkable divergence of spacer sets in poultry versus house finch arrays. We also detected further evidence for an ongoing inactivation of the CRISPR-Cas system in the house finch host over time. Second, we functionally characterized the CRISPR-Cas system of two *M. gallisepticum* strains infecting poultry (S6 strain) and house finch (CA06 strain) hosts. Using *in vitro* and *in vivo* approaches, we determined the PAM specificities of the MgCas9 and showed a correlation between a modification of the PAM signature, the evolution of the spacer repertoires, and the evolution of the PI domain of MgCas9. Such radical change in the CRISPR-Cas defence system of *M. gallisepticum* may have implications for the rapid adaptation of *M. gallisepticum* to its novel host (Bonneaud et al., 2018; Tardy et al., 2019).

## Results

### Comparison of the CRISPR-Cas locus across *M. gallisepticum* isolates

CRISPR-Cas sequences were extracted from previously published whole genome sequences of 19 poultry and 8 house finch strains (Leigh et al., 2019a, 2019b; Papazisi et al., 2003; Song et al., 2021; Szczepanek et al., 2010; Tulman et al., 2012), as well as from a further 79 newly-sequenced genomes of strains collected in house finches between 1994-2015 (Table S3). A single CRISPR locus was found in all *M. gallisepticum* isolates obtained from house finches (Figure 1A), with the exception of 6 isolates (1 collected in 2006, 3 in 2007, 1 in 2008 and 1 in 2015). In all 6 of these isolates, the CRISPR-Cas system was found to have been inactivated due to the loss of part of the locus harbouring the tracrRNA and the *cas* genes (i. e. *cas9, cas1, cas2* and *csn2*), leaving only the CRISPR array. An extended analysis of the genome region on both sides of the CRISPR locus further suggested this system being part of a larger defence island, also including a restriction-modification system, a toxin-antitoxin system and a putative S8 serine peptidase (SI-2). For at least the 2 isolates with a completely assembled genome (strains collected in 2006 and 2008), the deletion was extended upstream of the CRISPR locus, including the putative S8 serine peptidase and other adjacent genes. The inactivation of the CRISPR-Cas system could also be predicted for a further subset of isolates based on several mutations found in their MgCas9 gene. Indeed, we were able to predict three different truncated versions of MgCas9 (thereafter named V1, V2 and V3) (Figure 1B, table S3). The MgCas9 truncated form V1 form was observed only once in a 1998 house finch isolate; it is truncated at position 1498 in the REC domain as a result of a single base change that generates a TAG STOP codon instead of CAG Gln codon. The V2 form was detected in 7 isolates sampled in house finches in 2001, 2009 and 2015; it was predicted based on a frameshift at position 3445 of the CDS, leading to a protein truncated of the last 120 amino acids within the PAM Interacting domain (PI). Finally, the V3 form was identified in 29 isolates collected in house finches in 2003, as well as between 2011 to 2015; it is truncated at position 1018 of the CDS within the REC lobe (which exerts a key role for DNA cleavage) as a result of a single nucleotide mutation changing a GAA Glu codon into a TAA STOP codon.

**Figure 1.**
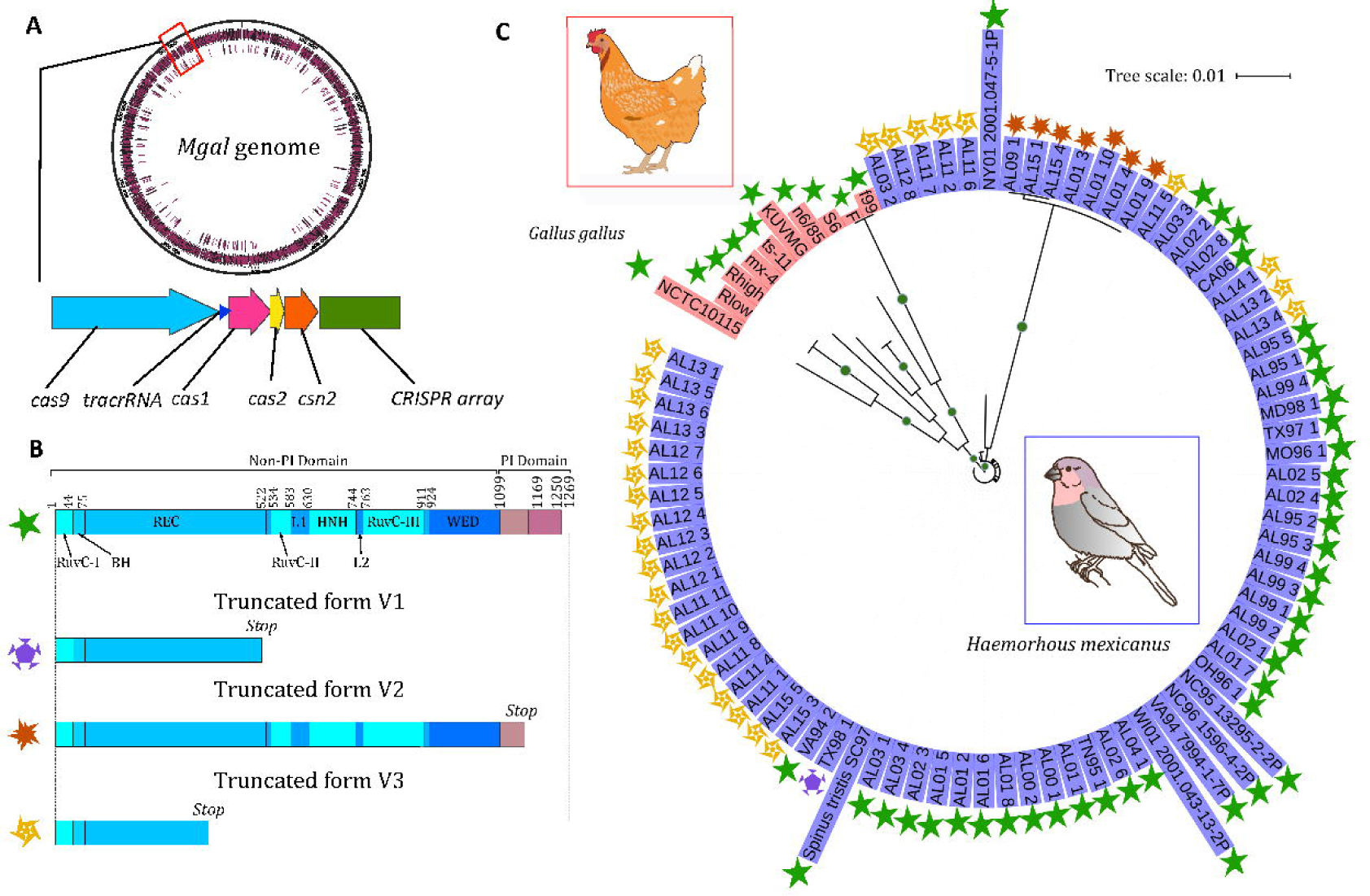
Presence and integrity of the CRISPR defence system in *M. gallisepticum* strains. **A -** Map of the *M. gallisepticum* genome highlighting the position of the CRISPR-Cas locus (red box) and its organization. **B** – Domains present in full-length and truncated forms of MgCas9 found in *M. gallisepticum* strains. Green star, strains with a complete CRISPR locus and a complete MgCas9 ; other motifs, inactivated CRISPR system with truncated forms V1, V2 and V3 of MgCas9. **C -** Phylogenetic tree of MgCas9. Strains isolated from poultry and house finch hosts are highlighted in red and purple, respectively. MgCas9 forms are indicated for all strains. Green dots on branches indicate bootstrap values >95%.

The pattern of occurrence of active and inactive MgCas9 in house finch isolates suggests an overlap of both forms over a relatively short period and followed by the complete inactivation of MgCas9. Indeed, all the house finch isolates for which we could predict a complete MgCas9 were collected between 1995 and 2006, whereas all the isolates collected after 2007 displayed either an inactivated MgCas9 or a partial deletion of the CRISPR locus. Isolates with active and inactive MgCas9 were therefore only found to co-occuring in house finches between 2001 and 2006. This gradual pattern of inactivation contrasts with what we see in poultry, as all strains collected in chicken and turkeys >60 years (∼1958-2019) are predicted to harbour an active CRISPR system and did not display any inactivating mutations in MgCas9.

### Analysis of spacers and identification of *M. gallisepticum* CRISPR targets

We compared the set of spacers inferred from the *M. gallisepticum* CRISPR locus across all studied *M. gallisepticum* isolates (Figure S1, Table S4), and compiled a dataset of 490 non-redundant spacers (Table S4C). As a result, we identified a total of 438 unique spacers from 19 poultry strains and 55 unique spacers from 85 house finch isolates (Table S4C). Across the *M. gallisepticum* poultry strains, we observed a wide diversity of spacers, with no spacer found conserved across all strains (Figure S1). Indeed, 66% of spacers from the 19 poultry strains were present in only 1-2 (0-10%) genomes, while 2% were found in 9-10 (40-50%) genomes. In contrast, we found a lower diversity of spacers across the CRISPR arrays of house finch isolates. A quarter (24%) of spacers found in house finches were shared across more than 80% of isolates, while over half (58%) were present in at least 40% of them, and 21 out of 55 (38%) spacers were present in 10% or less of house finch isolates.

We found only 3 spacers that were shared across poultry and house finch isolates (Table S4C). Spacer SP_516* was identified in the poultry vaccinal strain ts-11, in 7 derivates obtained by chemical mutagenesis (K6372, K6356, K6212B, K6216D, K6208B, K5322C, K2966) and in all but 10 of the house finch isolates. Spacer SP_119* was found in 4 ts-11 derivate strains (K6369, K6208B, K5322C and K6222B) and in 3 house finch isolates for which the CRISPR-Cas9 system was predicted to be inactive (169125442_2001, 13_09_02_NC and 2015_J). Finally, spacer SP_325* was shared between the poultry strains R_Low_ and R_High_ and by 5 house finch isolates (NC95, NY01, WI01, NC06 and NC08). When considering the pattern of spacer acquisition over time in house finches, we found that most new spacers (i.e. 46 out of 55; >80%) had been integrated before 2000 and were therefore present in the early years of *M. gallisepticum* emergence in house finches. Surprisingly, despite the inactivation of MgCas9 from 2007, we also found evidence for the acquisition of six new spacers between 2009 and 2015 (Table S4C).

We investigated the potential origin of *M. gallisepticum* spacers using Blast queries against PHASTER, GenBank and IMG databases (Tables S4D-S4G), as described in (Vink et al., 2021). Out of 490 spacers, 431 (88%) showed no hits in the explored databases. The PHASTER database returned hits on known bacterial phages for 34 spacers. GenBank returned hits on bacterial genomes (11 spacers) and phages (4 spacers). IMG returned hits on bacterial genomes for 8 spacers. In addition, two self-matching spacers were found in 2 strains of *M. gallisepticum* (ts-11 and NTCTC1015). Overall and across the 57 spacers with identified hits outside *M. gallisepticum* genomes, we noted 6 spacers that were present in house finch isolates, and 51 in those from poultry. Out of the 19 spacers with hits on bacterial genomes, 10 returned non-CRISPR genomic regions of mycoplasma. Among them, 6 spacers from poultry strains (S6, KUVMG001, 6/85, ts-11 and ts-11 derivates) matched to the genome of *Mycoplasma imitans,* a duck pathogen phylogenetically related to *M. gallisepticum* (Table S4F); these potential protospacers are localized in a *M. imitans* chromosomal region of 14 kb that is not found in the *M. gallisepticum* genomes used in this study, contrary to other surrounding loci. In this region, we identified several genes considered as essential for the mobilization and transfer of Mycoplasma Integrative and Conjugative Elements (MICE) (Figure S2) (Calcutt et al., 2002; Dordet Frisoni et al., 2013; Meygret et al., 2019). This suggested those 6 spacers might be directed against a potential mobile element of 14 kb from bird mycoplasma, although no trace of this element has been detected in the *M. gallisepticum* genomes studied here. Taken together, these results indicate that *M. gallisepticum* CRISPR-Cas9 systems are mostly directed against unknown targets and that predicted targets include most predominantly phages, as well as a newly identified ICE-like element of a bird mycoplasma.

### Diversity and evolution of Cas proteins

We undertook further *in silico* analyses to document the intraspecific diversity of *M. gallisepticum* Cas proteins across isolates displaying a complete functional locus. To do so, we aligned the protein sequences of MgCas9 and constructed a phylogenetic tree (Figure 1C). The isolates were clearly distributed into two main branches, with a clustering of the poultry MgCas9 sequences in one branch and of the house finch ones in another. House finch MgCas9 sequences were found to be >99% identical to each other and the copy present in the CA06 isolate was selected (as in (Gasiunas et al., 2020)) to be compared with a poultry MgCas9 from the S6 strain. This strain was chosen because it has not been mutagenized and it has been proven to be transformable, which was required for further experimental approaches. While being 96% identical in sequence, the highest proportion of amino acid variations was found within the PI domain, which is involved in the interaction with the PAM sequence (Figure 2A). Indeed, sequence variations of 8.7% and 3.1% were calculated for the PI domain and the other domains of the protein, respectively.

**Figure 2.**
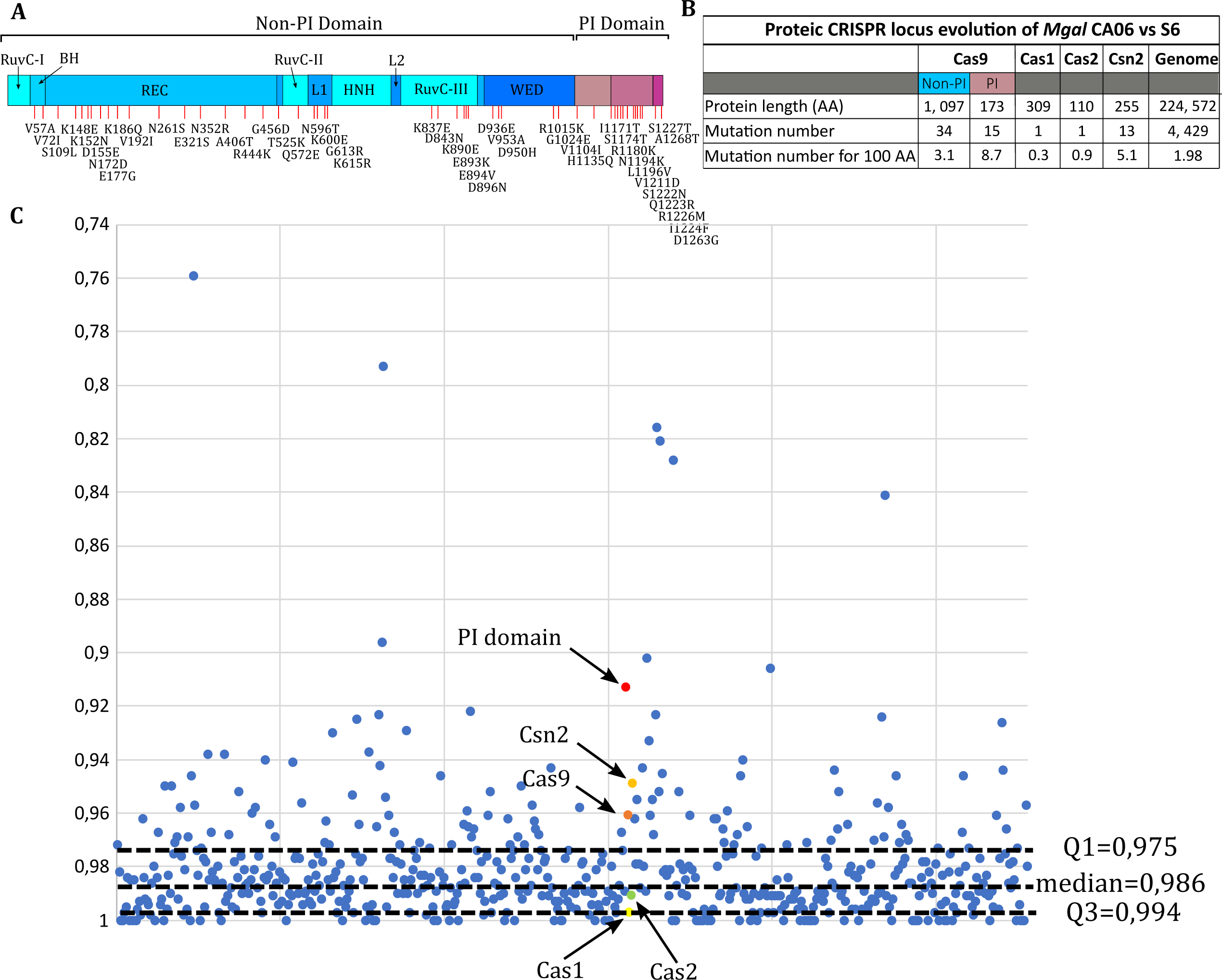
Evolution of the Cas proteins from *M. gallisepticum* poultry and house finch isolates. **A** - Representation of MgCas9 protein variable positions (red dash) between strains S6 (poultry) and CA06 (house finch) isolates. For each mutation, the corresponding amino-acid substitution is indicated. **B**- Protein sequence comparisons between CRISPR effectors of S6 and CA06 isolates. **C** - Protein identity plot. Percentages of identity are represented in the graph for each protein found in *M. gallisepticum* S6 genome and CA06 genome from PATRIC analysis.

When considering other Cas proteins encoded within the CRISPR locus, we found that Csn2 sequences exhibited more variation (5.1%) than either Cas1 (0.3%) or Cas2 (0.9%) (Figure 2B). A proteome-wide comparison of *M. gallisepticum* CA06 and S6 isolates showed that Cas9 and Csn2 are part of the 12% most divergent proteins between the two isolates (Figure 2C, Table S4). Together, our results suggest that the jump of *M. gallisepticum* from poultry into house finches has been marked by a noticeable evolution of the MgCas9 PI domain and of Csn2, both of which are key proteins in the ability of CRISPR-mediated immunity to acquire new spacers (Heler et al., 2015; Wei et al., 2015). Since the PI domain of Cas9 is involved in PAM recognition, our results further suggest that the jump might have been associated with a change in the PAM recognition specificity of MgCas9, a pattern which is consistent with a change in pathogen pressure experienced by *M. gallisepticum* within the novel finch host.

### *In vitro* PAM specificity differs between poultry (S6) and house finch (CA06) MgCas9

We used an *in vitro* assay previously developed (Gasiunas et al., 2020; Karvelis et al., 2019) to compare the PAM recognition preference of MgCas9 between the poultry strain S6 and the house finch isolate CA06. *In vitro* -produced MgCas9/sgRNA ribonucleoparticles were used for the cleavage of a library of plasmids containing a target sequence and a PAM sequence randomized on 7 positions (Figure 3A, Figure S3, Table S6). After three independent *in vitro* cleavage of the plasmid library by MgCas9 S6 or MgCas9 CA06, molecules containing a recognized PAM sequence were captured using DNA ligation of adapters, enriched by PCR amplification, and submitted for deep sequencing. Cleavage was observed three bases upstream from the PAM sequence at >99.95% and results obtained for replicates of the two series of were nearly identical (Figures S4A and S4B). Out of the 16,306 PAM sequences contained in the library, 10,928 and 9,709 PAM sequences were detected at least once after cleavage by MgCas9 S6 and MgCas9 CA06, respectively. The 1,000 most frequent PAM sequences were selected for each MgCas9 including 74 PAM sequences that were common to both MgCas9 S6 and MgCas9 CA06 top1,000 PAM sequences. Normalized frequencies were calculated, taking the bias of the original plasmid library into account (Table S6F); frequency matrices and sequence logos were produced (Figures 3B and 3C, Table S6G).

**Figure 3.**
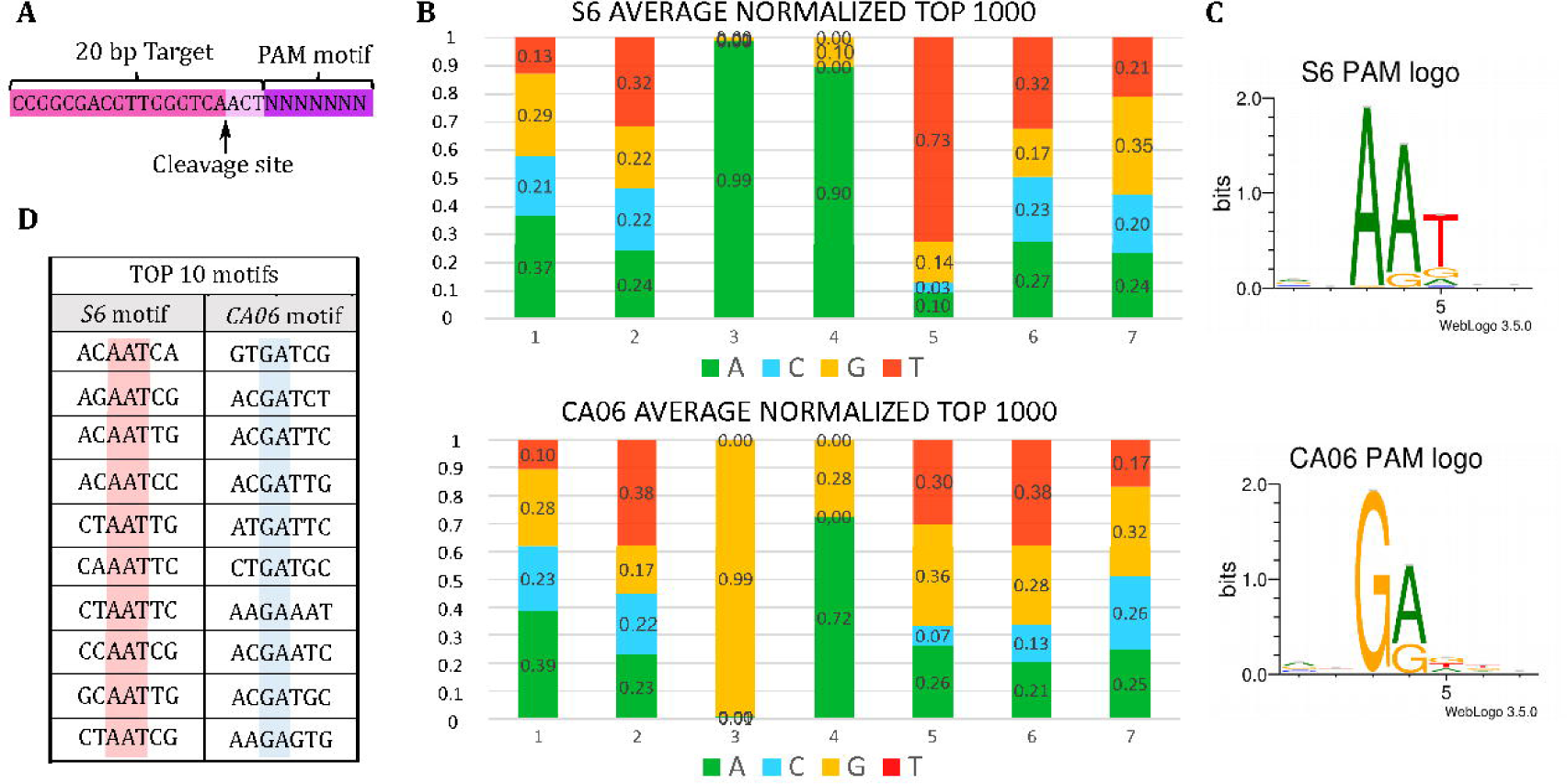
*In vitro* characterization of PAM sequences recognized by MgCas9 from poultry (S6) and house finch (CA06) strains. **A**- Position of the observed cleavage site for the two MgCas9.**B**- Position Frequency Matrix (PFM) observed after cleavage of a 7-positions PAM degenerated library by MgCas9 from S6 and CA06 *M. gallisepticum* strains. Frequencies were averaged on three replicates and corrected considering the bias in the distribution of PAM sequences in the initial plasmid library. Adenine are represented in green, Cytosine in blue, Guanine in yellow and Thymine in red. Frequencies of each base at each position are indicated on the graph. **C**- Web logos of consensus PAM sequences recognized by MgCas9 S6 or CA06. **D**- Top 10 of the best motifs recognized *in vitro* by MgCas9 S6 and CA06.

A comparison of the PAM recognition preference of MgCas9 S6 and MgCas9 CA06 revealed that the PAM sequences recognised by both correspond to a 5-letter motif, with a marked bias observed at positions 3, 4 and 5 only. Indeed, two lines of evidence indicate that the 5-position PAM sequences recognised by MgCas9 from poultry S6 and house finch CA06 strains differ *in vitro*. First, a sequence logo comparison revealed that PAM preferences of MgCas9 proteins were clearly different between the two isolates. The main difference was observed at position 3 with 99% of sequences displaying an A in MgCas9 S6 and a G in MgCas9 CA06 (Figures 3B). Position 4 was preferentially A or G in both cases, with a bias towards A more marked in MgCas9 S6. In position 5, a T was present in 73% of sequences retrieved after cleavage by MgCas9 S6, compared to 30% by MgCas9 CA06, suggesting a notable difference at this position between the two isolates. Second, we found that the top 10 PAM sequences most frequently recognized differed between S6 and CA06 MgCas9 (Figure 3D).

To confirm the difference of PAM recognition specificity, we selected two PAM sequences, specifically recognized by MgCas9 S6 (Motif 1, AT**AAA**AA) and MgCas9 CA06 (Motif 2, AA**GAG**AA), respectively. Three plasmids were constructed containing the target sequence followed by one of the two selected PAM sequences or by the 7 first nucleotides of Direct Repeat (DR) sequence interspacing spacers in *M. gallisepticum* CRISPR arrays (Motif 3, GT**TTT**AG) (Figure S5A). After *in vitro* cleavage of these plasmids by *in vitro* -produced MgCas9/sgRNA complexes, we observed that MgCas9 S6 efficiently cleaved Motif 1 but not Motif 2, whereas MgCas9 CA06 preferentially cleaved the Motif 2 (Figure S5B). By contrast, the control Motif 3 remained uncleaved after incubation with one MgCas9 or the other. This confirmed the difference of PAM specificity of MgCas9 originating from poultry and house finch strains, *in vitro*. Together, our results show that poultry and house finch MgCas9 recognise different 5-letter motifs in the PAM sequences and display different cleavage specificities of PAM sequences as a result.

### *In vivo* PAM specificity of MgCas9 is driven by PI domain evolution

We characterised the PAM specificity of poultry (S6) and house finch (CA06) MgCas9 using *in vitro* and *in vivo* approaches. Based on the results previously obtained *in vitro*, we selected four PAM motifs found to be differentially cut by poultry (S6) and house finch (CA06) MgCas9 (Figure 4A). Motifs S6-1 and S6-2 were found to be recognized by MgCas9 S6 *in vitro*, but poorly by MgCas9 CA06, while motif CA06-3 was cleaved by MgCas9 CA06 but not MgCas9 S6, and CA06-4 was efficiently recognized by MgCas9 CA06 but only weakly by MgCas9 S6. To investigate whether these findings obtained *in vitro* are representative of processes occurring *in vivo*, we conducted interference assays on both the wild-type *M. gallisepticum* S6 strain and on a genetically engineered *M. gallisepticum* S6 (thereafter named *M. gallisepticum* S6 mod PI CA06) in which the MgCas9 PI domain was seamlessly replaced by the homologous domain of the MgCas9 CA06 (Ipoutcha et al., 2022). The *in vivo* cleavage assay was based on the transformation of *M. gallisepticum* with a replicative plasmid containing the third spacer naturally found in the CRISPR array of strain S6, followed by a selected PAM candidate (Figure 4B). A plasmid that contained the Direct Repeat of the CRISPR array as a PAM sequence was used as a negative control.

**Figure 4.**
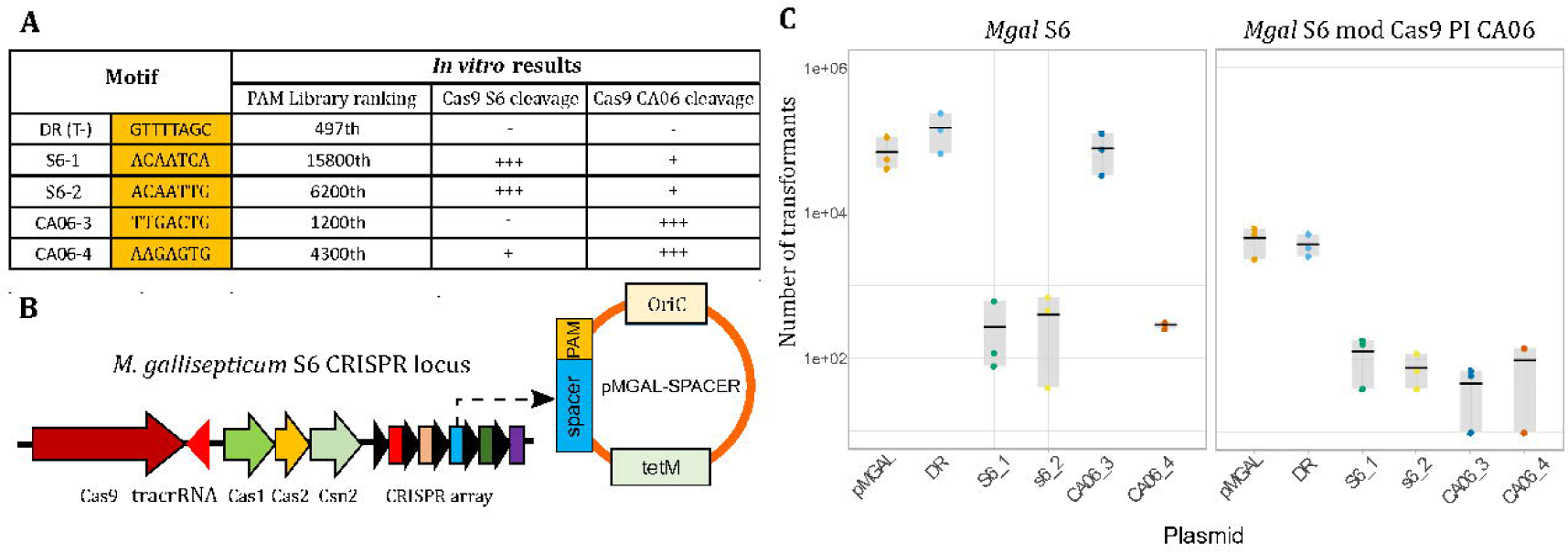
Evaluation of MgCas9 PAM specificity *in vivo* using a plasmid interference assay. **A**- PAM sequences for *in vivo* interference assays were selected from *in vitro* results. DR is the direct repeat sequence naturally present in the *M. gallisepticum* CRISPR array; this motif cannot be recognized by MgCas9 and is used as a positive control of transformation and negative control for cleavage. Four other PAM sequences where chosen, that are differentially cut *in vitro* by MgCas9 S6 and CA06. PAM sequences frequently retrieved after cleavage are noted “+++” whereas lowly and not recognized sequences are indicated by “+” and “-“, respectively. **B**- Plasmid design for interference assays in *M. gallisepticum* S6 and *M. gallisepticum* S6 mod PI CA06. Plasmid pMgal contains *M. gallisepticum oriC*, a *tet(M)* resistance marker and the third spacer which is present in the S6 CRISPR array, followed by selected PAM sequences. **C**- *In vivo* result of transformation assays for the two *M. gallisepticum* strains. The numbers of transformants obtained after three transformation replicates are represented as a dot plot.

Transformations with plasmids containing PAM sequences expected to be cut by MgCas9 S6 (S6-1 and S6-2) led to 100-1,000 fold less transformants compared to transformation with the control plasmid (Figure 4C). This confirmed that the CRISPR-Cas9 system is functional as a defense system in *M. gallisepticum* S6 and that PAM sequences recognized *in vitro* by MgCas9 S6 are also recognized *in vivo*. Similarly, plasmids harbouring PAMs frequently cleaved by MgCas9 CA06 *in vitro* were also cleaved by MgCas9 mod PI CA06 *in vivo*, demonstrating that the hybrid MgCas9 is fully active. Interestingly, in this interference assay, motifs that were poorly recognized *in vitro* (i.e. CA06-4 for MgCas9 S6; S6-1 and S6-2 for MgCas9 CA06), were efficiently cleaved *in vivo* (see SI-3 for further comparisons of *in vitro* and *in vivo* findings). Finally, as we expected, we found that the PAM CA06-3 was not recognized by MgCas9 S6, but was efficiently cleaved by the hybrid MgCas9 that included a CA06 PI domain. This result confirms that the specificity of MgCas9 is mediated by the PI domain and suggests that mutations in the MgCas9 of house finch isolates has led to a change in PAM specificity.

## Discussion

We studied the genomes of the bacterial pathogen, *M. gallisepticum*, that were collected in the original poultry host and from house finches periodically over a 21-year span following a host shift. Our focus was investigating the impact of a dramatic change in the host environment on the evolution of the pathogen’s CRISPR-Cas defence system. First, our results were consistent with a previous observation, made from a small number of isolates, that there has been a gradual inactivation of MgCas9 in the house finch host since colonization. Indeed, we found that, while all poultry strains displayed a an active CRISPR-Cas system, both active and inactive forms of MgCas9 were present in house finch isolates collected in the initial 12 years following the host shift. After the twelfth year following colonization, however, all *M. gallisepticum* collected from house finches displayed either an inactivated MgCas9 or a partial loss of their CRISPR locus (*cas* genes). Second, of the 490 unique spacer sequences detected across poultry and house finch isolates, only three were shared across both avian hosts, with poultry isolates displaying a greater diversity of sequences than house finch isolates. Third, we found that the protein sequences of MgCas9 of poultry and house finch isolates clustered into two distinct phylogenetic branches, with very little variation detected among house finch MgCas9 protein sequences. Within MgCas9, the PAM-interacting (PI) domain (involved in the specific interaction between Cas9 and the protospacer adjacent motif (PAM) that follows the target DNA sequence) and the Csn2 protein sequences were found to be most variable across poultry and house finch isolates. Fourth, we found that MgCas9 recognises a 5-letter motif in the PAM sequence. The motif recognised differed between poultry and house finch MgCas9 and, accordingly, we measured variation in cleavage efficiencies *in vitro*. The poultry MgCas9 was more effective at cleaving its target PAM sequences than the house finch MgCas9, while the house finch MgCas9 was most effective at cleaving its target PAM. Finally, our results confirmed that differences in the PAM specificity of MgCas9 are driven by differences in the PI domain between poultry and house finch strains. Indeed, differences in specificities were replicated across a wild-type poultry strain (S6) and a genetically engineered mutant of that strain in which the PI domain of MgCas9 was replaced by the homologous domain of a house finch isolate (CA06), with the latter more effective at cleaving PAMs from house finch isolates. Together, these results suggest that significant evolutionary changes took place in *M. gallisepticum*’s CRISPR-Cas defence system following the jump into the novel house finch host. We documented an initial pattern of acquisition and loss of CRISPR spacers, and of changes in PAM specificity driven by the MgCas9 PI domain. This observation is consistent with an evolutionary response to exposure to a new community of phages and MGEs. That such initial functional changes were followed by the subsequent inactivation of this defence system, however, highlights the complexity of the selection pressures that it faced.

Our study on the CRISPR-Cas9 system of *M. gallisepticum* isolates collected in both poultry and house finch hosts provides support for previous work, while also broadening our understanding of the complexity of this system across vertebrate host environments. First, mycoplasma CRISPR-Cas9 systems have previously been classified as Type II *in silico*, which include the typical set of *cas* genes, *cas1*, *cas2*, *cas9* and *csn2* (Ipoutcha et al., 2019). Accordingly, we found that the CRISPR-Cas systems of *M. gallisepticum* poultry S6 and house finch CA06 isolates are typical of the Type II systems. Second, our results are also consistent with a previous study that undertook the first functional investigation of the Cas9 endonuclease in a mycoplasma system as part of a larger initiative to characterize 79 Cas9 orthologs representative of 10 major clades in a Cas9 evolutionary tree (Gasiunas et al., 2020). In this study, the house finch isolate CA06 was selected as representative mycoplasma Cas9 with a 5-positions motif NNGAD as its consensus PAM (Gasiunas et al., 2020). Our results provide confirmation for MgCas9 CA06’s consensus PAM. Interestingly, however, we identified a different PAM motif (NNAAT) in the MgCas9 of the poultry strain S6, which was found to exhibit an ultra-dominant A at position 3 (99%) and a marked trend (73%) towards a T at position 5. In addition, we found that this difference in the PAM recognition preference of MgCas9 was associated with differences in both the composition and diversity of the spacers between poultry and house finch isolates, suggestive of a complete reset of the CRISPR array following the host shift.

An active adaptive immunity system like CRISPR-Cas is expected to be maintained only when the benefits of such a system outweigh the costs of sustaining it (Dimitriu et al., 2020; Koonin, 2019; Westra & Levin, 2020). The main benefit of an active CRISPR-Cas system is the protection that it confers against phages and MGEs. For example, in the case of *M. gallisepticum*, the presence in poultry strains of 6 spacers matching to the genome of the duck pathogen *M. imitans* suggests that avian mycoplasma bacteria are under selection to actively protect themselves against invasion by circulating MICEs. However, maintaining an active CRISPR-Cas undoubtedly incurs fitness costs. Such costs can arise, for instance, when CRISPR-Cas generates metabolic costs (Hall et al., 2021; Vale et al., 2015) and genetic conflicts (Hall et al., 2021; Vale et al., 2015) or when it is less effective at preventing infections than other defence processes (Stern et al. 2010; Broniewski et al. 2020; Watson et al. 2024).CRISPR-Cas can also create a risk of self-targeting spacers (Stern et al., 2010; Wimmer & Beisel, 2020), and can prevent the acquisition of fitness-enhancing genes through horizontal gene transfer (HGT) (Alduhaidhawi et al., 2022; Bikard et al., 2012; Dimitriu et al., 2020; Kogay et al., 2024; Levin, 2010; Mackow et al., 2019; Palmer & Gilmore, 2010; Pursey et al., 2022; Wheatley & MacLean, 2021). The balance between the costs and benefits of maintaining an active CRISPR-Cas will therefore depend on CRISPR-Cas’ contribution to the bacterium’s global defence process, on its non-defence roles, and on the specific selective pressures exerted by different host environments.

That there were isolates in circulation with an active (albeit reconfigured) CRISPR-Cas system in the 12 years following the jump into house finches indicates that there were benefits to the maintenance of this defence mechanism during that time period (Figure 5). The retention of an active CRISPR-Cas system is particularly striking when one considers the function of the system was maintained even with a large number of mutations (49 amino-acid changes when comparing MgCas9 S6 and MgCAs9 CA06) that accumulated in the house finch *cas9* gene following the jump. The reasons for such benefits are likely to be found in the distinct PAM specificity of MgCas9 in house finch isolates, alongside salient patterns of loss of ‘poultry-type’ spacers and acquisition of new spacers elements. Indeed, spacers that are no longer in use are expected to be lost, while new ones that confer resistance against novel pathogenic targets should be gained. *M. gallisepticum* isolates from house finches display dramatic changes both in spacer set and MgCas9 PAM specificity, which suggest that the bacterium was faced with a massively different community of phages and invasive MGEs in the novel host. Indeed, such a finding is consistent with previous evidence of a rapid evolution of PAM specificities reported to occur in response to an escape from detection by phages and MGEs displaying mutations in their PAM motifs (Deveau et al., 2008; Edraki et al., 2019; Paez-Espino et al., 2015). Whether *M. gallisepticum* experienced novel pathogen-driven selection in house finches, and the role that this might have played in facilitating or hindering its emergence remain to be determined. Regardless, our results give rise to the hypothesis that the successful emergence of a bacterial pathogen in a new host species will not only depend on its ability to establish and spread in the new host, but also on its ability to cope with a new landscape of phages and other invading DNAs.

**Figure 5.**
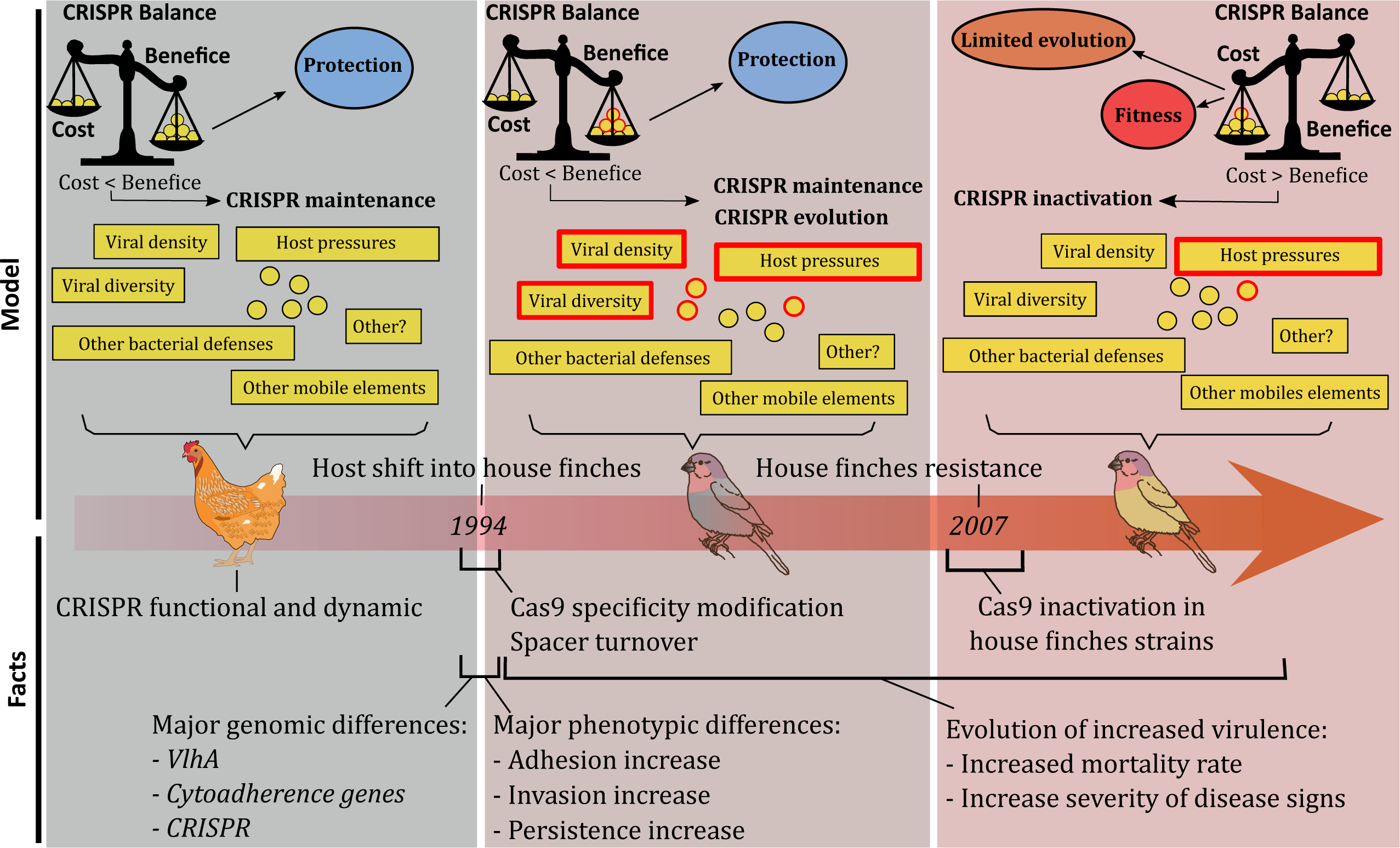
Hypothetical model of evolution of the CRISPR-Cas system in *M. gallisepticum* as it jumped from poultry and emerged into the novel house finch host. Phenotypic and genetic changes accompanying each step are obtained from the literature.

As *M. gallisepticum* adapted to its new host and novel sets of phages and invasive MGEs, the costs of maintaining an active CRISPR-Cas system seem to have outweighed the benefits of such a system. Isolates displaying an inactive CRISPR-Cas system were also found in circulation over the 12 years following the jump into house finches and, by 2007, all the isolates collected either displayed an inactivated MgCas9 or the complete loss of their *cas* genes locus, leaving only the CRISPR array (Figure 5). Indeed, *cas9* inactivation evolved at least three times over that period from different point mutations leading to stop codons. Across bacteria, the switching off of the CRISPR-Cas systems has been proposed as a mechanism by which to respond to selection (Patterson et al., 2017). In *Escherichia coli*, for example, CRISPR-Cas is thought to restrict adaptive change by limiting the acquisition of potentially beneficial genes through horizontal transfer, giving rise to a trade-off between defence and adaptive potential (García-Martínez et al., 2018). As a result, we might expect the balance of costs and benefits of maintaining an active CRISPR-Cas system to have shifted for *M. gallisepticum* in the face of a significant change in selection in the novel host.

One major selective event faced by *M. gallisepticum* after the jump into house finches was the evolution of host resistance to infection (Bonneaud et al. 2011; Bonneaud et al. 2019; Dowling et al. 2020). The jump gave rise to an epidemic that spread quickly and is thought to have killed millions of house finches (Dhondt et al., 1998; Nolan et al., 1998). The high mortality rates of *M. gallisepticum* in house finches was a result of *M. gallisepticum* localizing in the mucosal surfaces of the conjunctiva and upper respiratory tract, causing a severe conjunctivitis that can lead to death in the wild through blindness-induced starvation or predation (Adelman et al., 2017; Roberts et al., 2001). The intensity of this selective event led to the spread of resistance in house finch populations from standing genetic variation and within only 12 years of epidemic outbreak (Bonneaud et al., 2011, 2018). As a result, by 2007, the proportion of resistant individuals in exposed house finch populations had risen to ∼80% (Bonneaud et al., 2011). Such host resistance was, in turn, found to have significant selective consequences for *M. gallisepticum*. Indeed, resistance favoured the evolution of increasing pathogen virulence through antagonistic coevolution (Bonneaud et al., 2018), with isolates causing ever greater host mortality and more severe signs of conjunctivitis over time (Tardy et al., 2019), as well as transmitting faster to uninfected sentinels (Bonneaud et al., 2020). The concordance in the timing of the complete inactivation of CRISPR-Cas (i.e., from 2007) and the need for evolutionary changes in virulence in response to widespread house finch resistance is striking. Further work is now required to test whether significant phenotypic and genetic changes have, indeed, taken place following the loss of a functioning CRISPR-Cas, as predicted if this defence system hindered the necessary adaptive responses.

In conclusion, our study of the CRISPR-Cas defence system of bacterial isolates collected from the original host, as well as from a novel host into which it jumped recently, reveals marked differences between both in term of: (1) the MgCas9 PAM recognition specificity, (2) the composition and diversity of the spacer repertoire, and (3) the gradual inactivation of CRISPR-Cas over time in the novel host. While the changes in MgCas9 PAM specificity and in the composition and diversity of spacers indicates a change in the landscape of phages and MGEs faced by the bacterium in the novel host, the subsequent inactivation of CRISPR-Cas is consistent with another important selection event. We hypothesize that the gradual inactivation of the CRISPR-Cas defence system, which took place over the first 12 years of the epidemic until this defence system became fully inactivated from 2007, occurred in response to the evolution of host resistance, which became widespread by 2007 and pushed for adaptive changes in the bacterium. Such adaptive changes would have necessitated the loss of a functional CRISPR-Cas defence system to take place. Together, our results highlight the need to consider not only the host-driven selection pressures a bacterium experiences, but also the complex interplay between phages and defence systems for better understanding the key factors driving the emergence of a pathogenic bacterium in a novel host.

## Material & Methods

### Oligonucleotides and plasmids

All oligonucleotides used in this study were supplied by Eurogentec and are described in Table S1. All plasmids constructed and used in this study are listed in Table S2. Detailed protocols for plasmid construction and other methods are provided in SI-1.

### Bacterial strains, culture conditions

*M. gallisepticum S6* (Tax ID: 1006581) was cultivated in Hayflick modified medium (Freundt, 1983) under a 5 % CO_2_ atmosphere. Tetracycline 5 µg/mL was used in selective conditions. *M. gallisepticum* S6 strain with a modified PI domain of MgCas9 was obtained as described in (Ipoutcha et al., 2022).

#### *In vitro* determination of MgCas9 PAM specificity

A 7N degenerated PAM plasmid library was provided by Dr. Gasiunas from Caszyme (Gasiunas et al., 2020). Randomness of the 7 positions of the randomized PAM was validated as described in (Karvelis et al., 2019). *In vitro* cleavage assays were performed as explained in (Gasiunas et al., 2020). Details of the protocol are provided in SI-1.

### Transformation of *M. gallisepticum*

*M. gallisepticum* cells were grown for 36 h before transformation. At pH 6.2 to 6.5, aliquots of 10 mL were centrifuged during 15 min at 6000 x *g*, 10°C. Cells were then resuspended with 5 mL of HBSS 1X wash buffer (ThermoFisher, 14065056) and centrifuged 15 min at 6000 x *g*, 10°C. The pellet was resuspended in 250 µL of CaCl_2_ 0.1 M and incubated for 30 min on ice. Cold CaCl_2_-incubated cells were gently mixed with plasmid DNA (10 µg) and 10 µg of yeast tRNA (ThermoFisher, AM7119). Then, 2 mL of 40 % polyethylene glycol 6000 (PEG) (Sigma, 11130) dissolved in HBSS 1X buffer were added to the cells. After 2 min of incubation at room temperature, contact with the PEG was stopped by addition of 20 ml of HBSS 1X wash buffer. The mixture was centrifuged 15 min at 6000 x *g*, 10°C and the cells were resuspended in 1 mL of modified Hayflick medium pre-warmed at 37°C. After 2 hours of incubation at 37°C, cells were plated on Hayflick selective plates. After 10-15 days at 37°C with 5 % CO_2_, colonies were counted, picked and re-suspended in 1 mL of Hayflick with selection for three passages (one passage ∼48 h).

## Acknowledgements

We are very thankful to Dr. Gasiunas from Caszyme for sharing the 7N PAM-degenerated pTZ57-7n library, the pET28ID116 plasmid encoding *M. gallisepticum* CA06 Cas9 and for appreciated advices on the protocol used for *in vitro* cleavage. We thank M. Staley for growing and shipping the house finch isolates and Andrea J. Dowling for growing the isolates, extracting their DNA and organising the sequencing of the genomes. We also want to thank Patrice Mora for helping in the constitution of the *M. gallisepticum* spacers database, Mammadou Sall for his contribution to the study of spacers and Alain Blanchard for much appreciated discussions on the work and manuscript.

## Author contributions

Thomas Ipoutcha: Conceptualization, methodology, validation, formal analysis, investigation, writing - original draft, writing - review & editing

Iason Tsarmpopoulos: methodology, investigation

Géraldine Gourgues: investigation

Vincent Baby: software, formal analysis, investigation

Paul Dubos: software

Geoffrey E. Hill: review & editing

Yonathan Arfi: Conceptualization, writing - review & editing, supervision

Carole Lartigue: Conceptualization, methodology, writing - review & editing, supervision

Patricia Thébault: methodology, software, formal analysis, investigation, data curation Camille Bonneaud: Conceptualization, writing - original draft, writing - review & editing

Pascal Sirand-Pugnet: Conceptualization, methodology, investigation, data curation, writing - original draft, writing - review & editing, supervision

