## Supplementary Information for "Evolution of the CRISPR-Cas9 defence system in *Mycoplasma gallisepticum* following colonization of a novel bird host"

**Supplementary Figures**

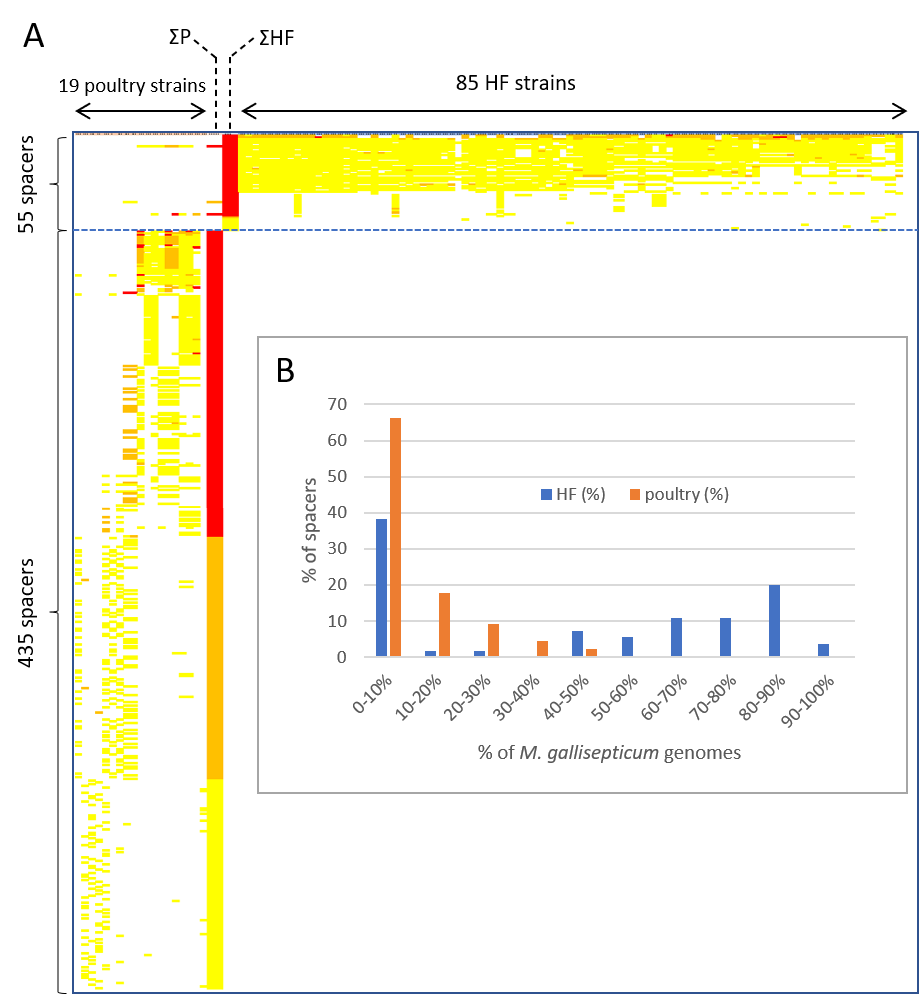

**Figure S1. Spacer distribution among *M. gallisepticum* genomes.** A. Overview of spacer occurrence among 19 poultry strains and 85 house finch strains. Each line corresponds to a spacer, each raw correspond to a strain. Spacer occurrence is indicated as follows: yellow, orange, red indicate spacers present on time, two times, more than two times in the genome; respectively. ΣP, sum of spacer occurrence among poultry genomes; ΣHF, sum of spacer occurrence among house finch genomes (same color code). B. Distribution of house finch spacers (blue bars) and poultry spacers (orange bars) among house finch and poultry *M. gallisepticum* strains.

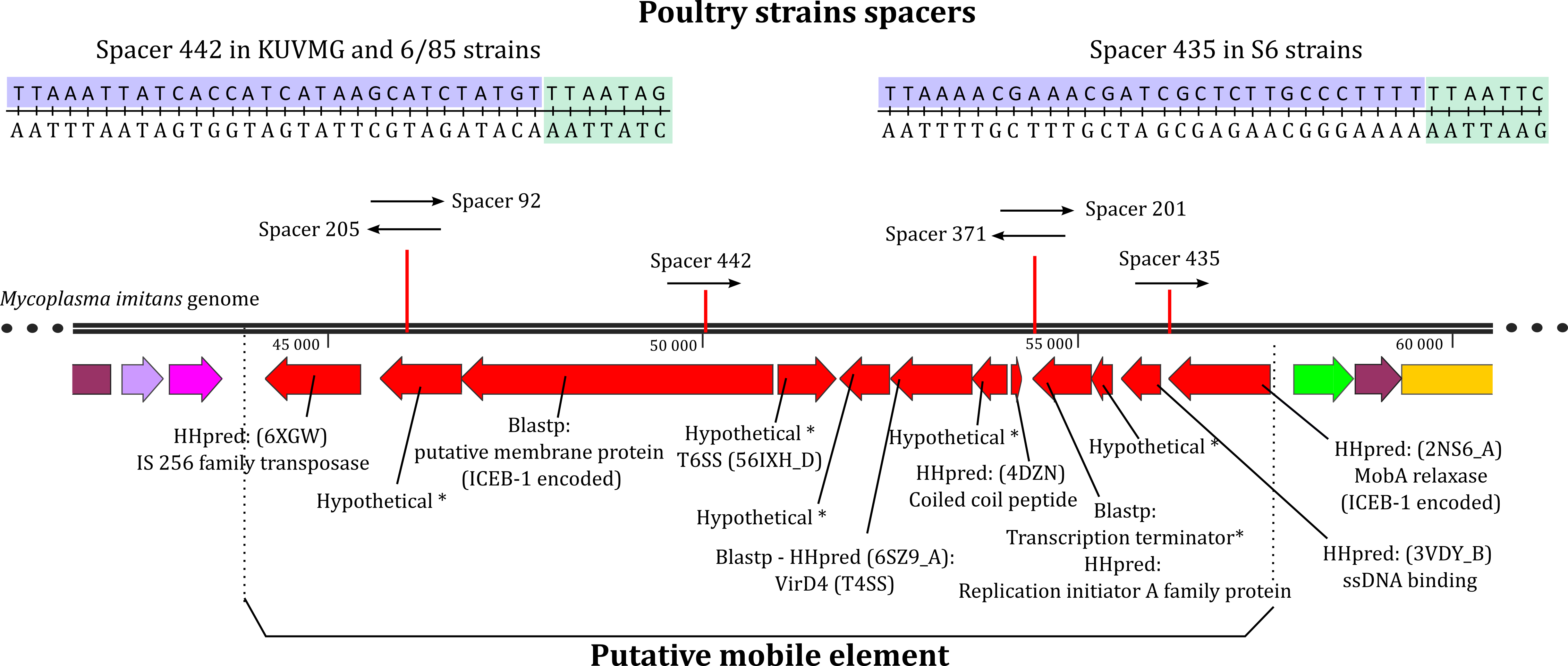

**Figure S2. Putative ICE-like element identified in *Mycoplasma imitans*.** The *Mycoplasma imitans* genome region encoding a putative ICE-like element and targeted by 6 spacers found in *M. gallisepticum* CRISPR arrays is represented. Genes in red were not found in *M. gallisepticum* genomes. Putative protein functions were proposed from BLASTp and HHpred searches. We also represent examples of two spacers (442 and 435) matches (light purple) and possible PAM sequence (light green).

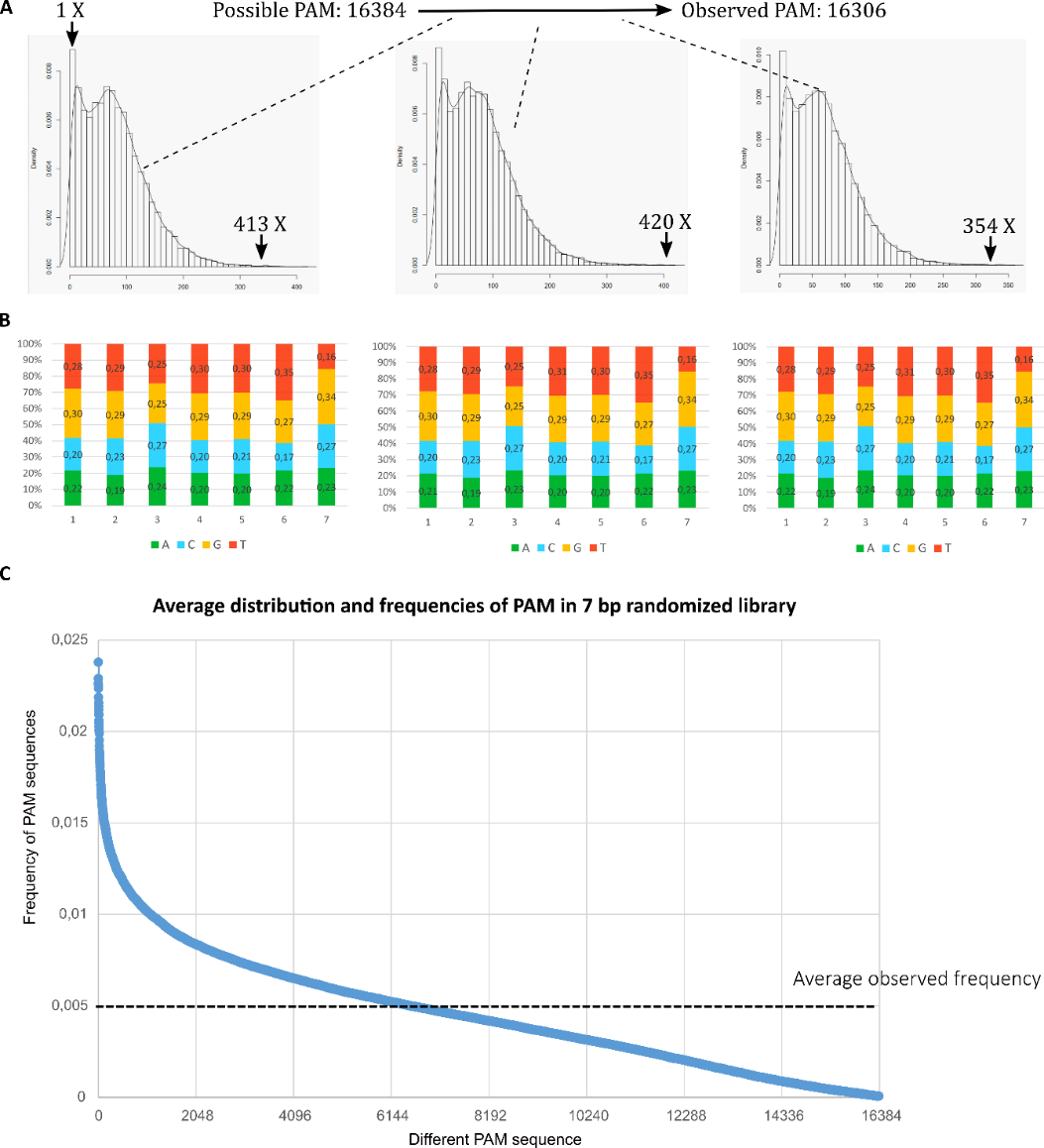

**Figure S3. Plasmid library validation. A-** Histograms of PAM motif distributions for three replicates are represented with minimum and maximum coverage. **B-** Position frequency matrix of the 7 bp randomized for each replicate. **C-** Average frequency of each PAM sequence observed in at least one of the three replicates was calculated. Average frequencies of PAM sequences in the 7 bp randomized library were ranked and represented as a graph. Average frequency observed was indicated by a dotted line.

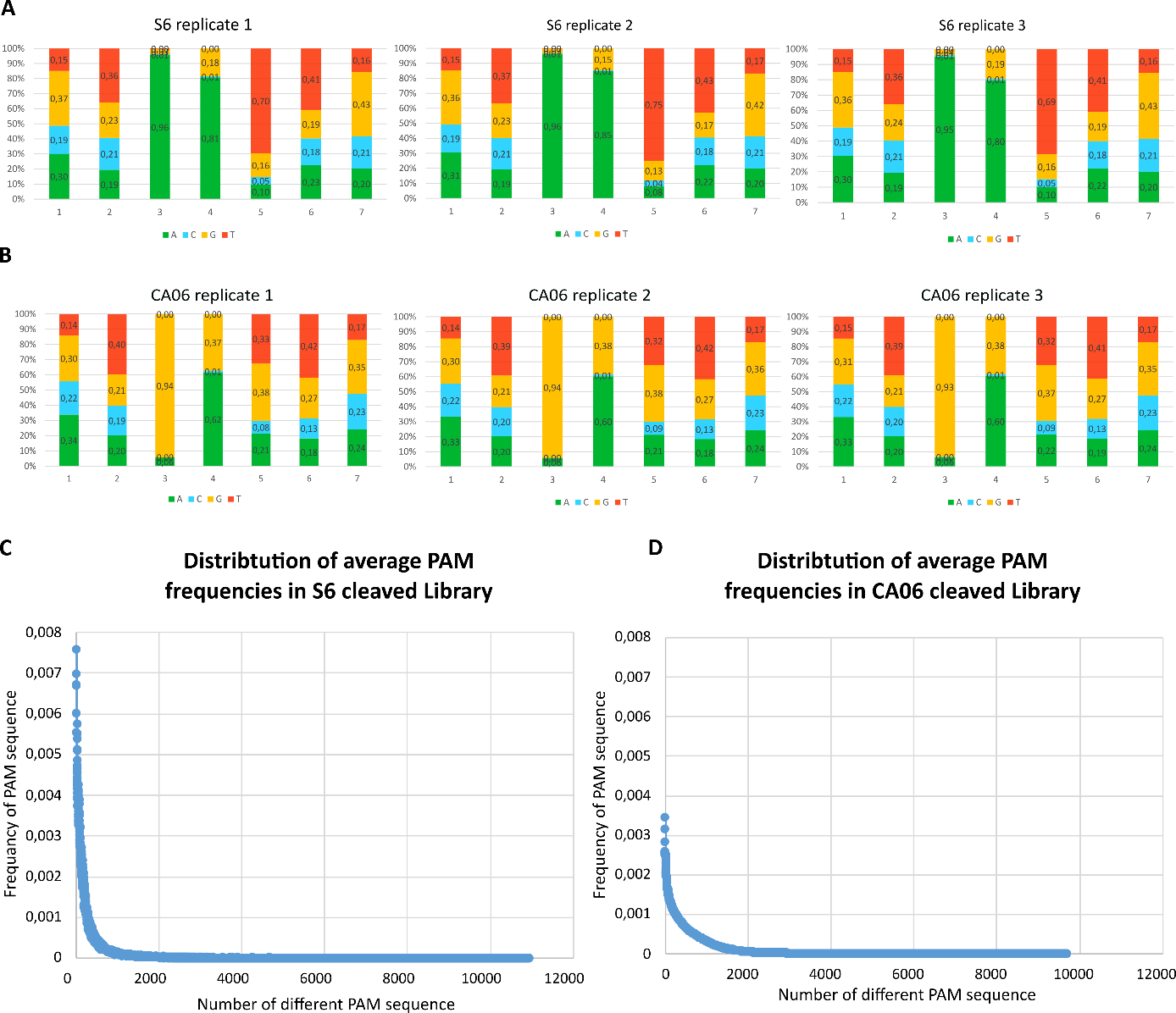

**Figure S4. PAM analyses after cleavage with MgCas9 from S6 (A) and CA06 (B) strains. A-** Position frequency Matrix of PAM cleaved sequence by S6 Cas9 for each replicate. **B-** Position frequency Matrix of PAM cleaved sequence by CA06 Cas9 for each replicate. **C-** Average frequency of each PAM sequence observed in at least one of the three replicates was calculated. Average frequencies were ranked and represented as a graph.

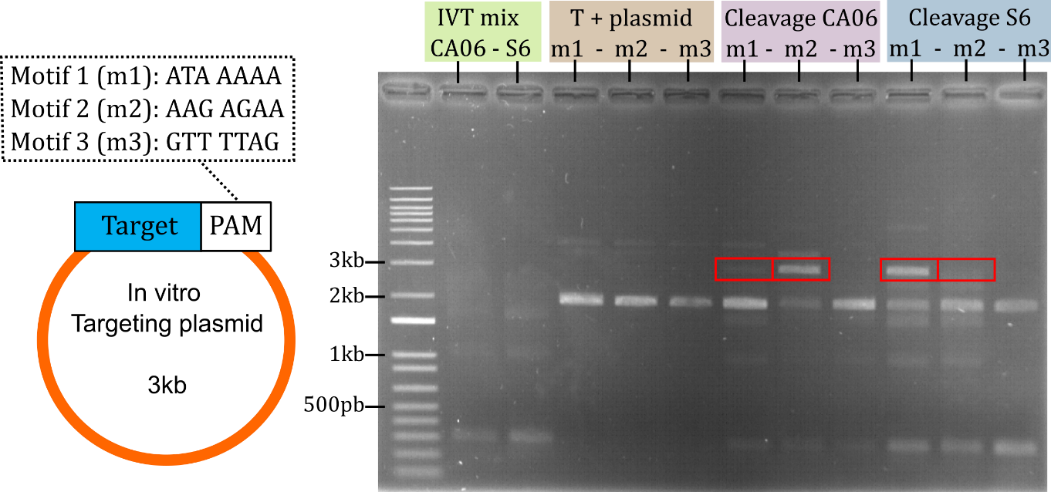

**Figure S5. *In vitro* plasmid cleavage assay with selected PAM sequences.** **A**- Design of the plasmids used for *in vitro* cleavage assay. PAM sequences m1, m2 and m3 were inserted downstream of a spacer targeted by the sgRNA to generate plasmids pTZ57-PAM_1, pTZ57-PAM_2 and pTZ57-PAM_3, respectively. **B**- Result of the *In vitro* cleavage after incubation or the plasmids with MgCas9/sgRNA complexes produced by IVT. Uncleaved plasmids were used as control (T+). Reaction products of IVT were also loaded on gel as a control of background bands.

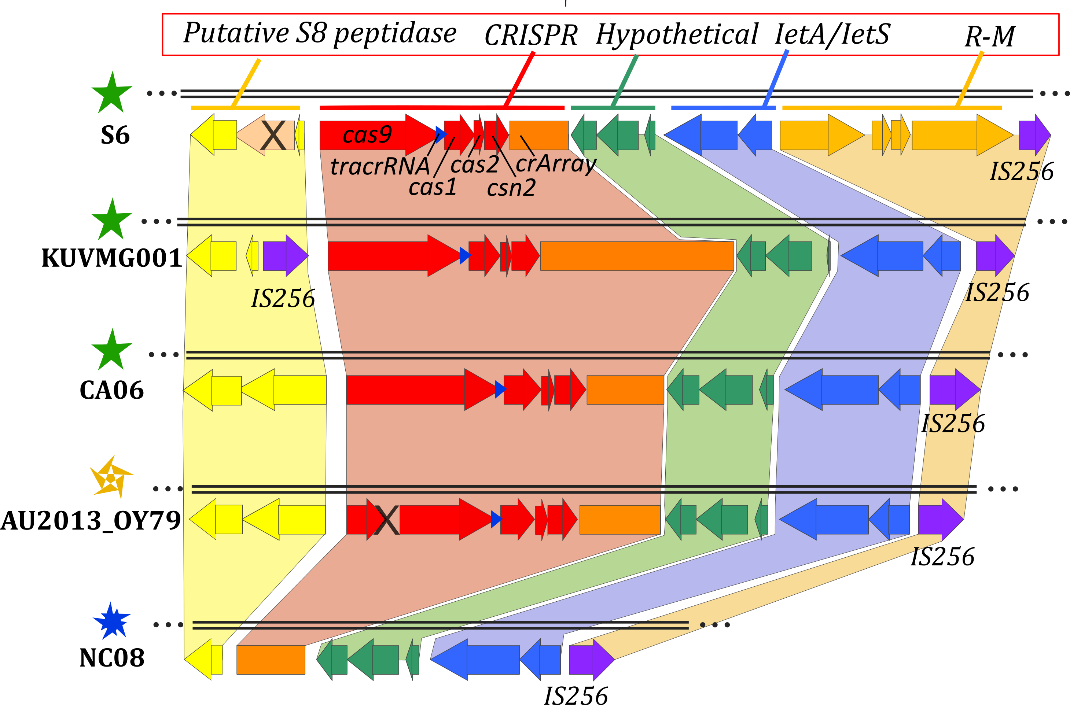

**Figure S6. Putative defence island in *M. gallisepticum* genomes*.*** Representation of the putative defence island in five different strains of *M. gallisepticum*. CRISPR system, IetA/IetS and Restriction-Modification system (R-M) were predicted using PADLOC tool (<https://padloc.otago.ac.nz>). Systems in green and yellow were found using manual inspection. An IS256 gene (purple) was also found at different locations in several strains. Star symbols refer to the integrity of the CRISPR locus (see Figure 1). Blue star symbol for isolate NC08 corresponds to strains with a complete deletion of *cas* genes.

**Supplementary text**

**SI-1. Materials and Methods**

**Genome sequences.** The genome of 79 *M. gallisepticum* isolates collected in house finch from 1994 to 2015 were sequenced using Illumina technology and draft assemblies were produced by C. Bonneaud et al. as part of a global comparative genomic study (unpublished). Other genomes used in this study were retrieved from GenBank and their accession numbers are available in Table S3. The CRISPR-Cas loci including *cas* genes, tracrRNA and arrays of spacers were extracted from the *M. gallisepticum* genomes for further comparative analyses. Their sequences are available in SI-5.

**Phylogenetic tree of MgCas9 and comparison of MgCas9 from S6 and CA06 strains.** Phylogenetic analysis of MgCas9 was performed using protein sequences and the phylogeny pipeline available at <http://www.phylogeny.fr/>. This pipeline includes the softwares Muscle, Gblocks, PhyML and TreeDyn. Representation of the tree was designed on iTOL (<https://itol.embl.de/>). For the truncated forms of MgCas9, the different parts of the ORF were artificially merged to include them in the tree. Protein alignments were analyzed on MEGA-X using Muscle alignment software (Kumar et al., 2018). Mutations rates between S6 or CA06 were calculated from SNP detected using whole genome nucleotide alignment in MAUVE (Darling et al., 2004). For mutation rates of proteins, concatenated protein sequences of whole genome were compare using Blast-p.

**Whole proteome comparison of S6 and CA06 strains.** Proteome comparison between S6 and CA06 strains was performed using the PATRIC tool (<https://www.bv-brc.org/app/SeqComparison> ) and fasta protein files from NCBI. Setting used: Minimum % coverage: 50%, BLAST E value: 1e-5, Minimum % Identity: 50%.

***In silico* analyses of *M. gallisepticum*** **spacers.** For each *M. gallisepticum* genome, spacers were detected using CRISPRfinder (<https://crispr.i2bc.paris-saclay.fr/Server/>) (database version: 2017-05-09). Newly sequenced House Finch strains draft assemblies obtained from Illumina short reads could contain uncertain nucleotides N. All spacers containing N tracks were removed to avoid an artificial increase of the spacer diversity. Using a homemade script, a matrix representing the occurrence of spacers in the *M. gallisepticum* genomes was produced (Table S4A). The 534 non-redundant spacers were next clustered by coverage and identity (>= 90% each) using CD-HIT (Li & Godzik, 2006), and a new simplified matrix representing the occurence of the 490 merged spacers was produced (Table S4C). PHASTER (update: Mar 29, 2018) (Arndt et al., 2016), Genbank, IMG (Chen et al., 2019) database were next requested to identify putative targets (Blastn >= 90% for identity and coverage) (Tables S4D, S4E, S4F). Spacers were also used in Blastn queries against *M. gallisepticum* genomes, spacers with more than one hit were also listed (Table S4-G).

**Construction of a minimal *M. gallisepticum*** **sgRNA.** Sequences of DR and tracrRNA were concatenated and the secondary structures of the hybrids were simulated using the mfold software at http://unafold.rna.albany.edu/. The sgRNA was designed according to (Jinek et al., 2012). The chimeric molecule was synthesized by IDT DNA company and further cloned in the plasmid pET28.

**Plasmids Constructions**

Plasmids pTZ57-Motif1, pTZ57-Motif2 and pTZ57-Motif3. The pTZ57-7n library vector provided by CasZyme (Gasiunas et al., 2020) was used as PCR template. Primers D17/D20, D18/D20 and D19/D20 and the Q5 High-Fidelity DNA Polymerase kit (NEB, M0491) were respectively used to amplify linear DNA fragment for pTZ57-Motif1, pTZ57-Motif2 and pTZ57-Motif3. PCR products were submitted to restriction with the DpnI enzyme (NEB, R0176S) following the manufacturer’s protocol. Both products were purified using GFX™ PCR DNA or Gel Band Purification Kit (GE-healthcare). The NEBuilder HiFi DNA Assembly Cloning Kit (NEB, E5520S) was used to assemble the fragments into the pTZ57-Motif1, pTZ57-Motif2 and pTZ57-Motif3 plasmids*.* Two µL of the assembly was transformed into *E. coli* NEB 5-alpha (NEB, C2987H). Transformants were screened after DNA minipreps using NucleoSpin Plasmid kit (Macherey-nagel, 740588.50) and enzymatic digestion. Sanger sequencing was performed for final verification.

Plasmids pET28-MgalS6Cas9 and pET28-sgRNA. The pET28-MgalCA06-Cas9 vector derived provided by CasZyme (Gasiunas et al., 2020) was used as PCR template. Primers B7/B8, A3/A4, A1/A2 B5/B6 the Q5 High-Fidelity DNA Polymerase kit (NEB, M0491) were respectively used to amplify MgalS6Cas9, sgRNA and backbone fragments. Cloning was performed using the same procedure as for pTZ57-Motif1 (see above) and resulted in the plasmids pET28-MgalS6Cas9 and pET28-sgRNA.

Plasmids pMGAL, pMGAL-Spacer-DR, pMGAL-Spacer-S6-1, pMGAL-Spacer-S6-2, pMGAL-Spacer-CA06-3, pMGAL-Spacer-CA06-4. The backbone shuttle vector pSRT2 was used. The origin of replication of *M. gallisepticum* has been identified by (Lee et al., 2008) as the area between MGA_0618 and MGA_0619. Primers E21/E22 were used and the Q5 High-Fidelity DNA Polymerase (M0491) kit to amplify the desired region. PCR product and pSRT2 vector were digested with the BamHI enzyme, following the suggested protocol. Both products were ligated following the T4 Ligase of Promega’s protocol thus creating the pMGAL plasmid. The construction was completed with an introduction through an XmaI digestion and Ligation of annealed Primers E23/E24 carrying the third spacer sequence from the *M. gallisepticum* S6 CRISPR locus. Primers E25/E26, E27/E28, E29/E30, E31/E32, E33/E34 and the Q5 High-Fidelity DNA Polymerase kit (NEB, M0491) were respectively used to change nucleotide sequence after target sequence and insert desired PAM sequence to construct pMGAL-Spacer-DR, pMGAL-Spacer-S6-1, pMGAL-Spacer-S6-2, pMGAL-Spacer-CA06-3, pMGAL-Spacer-CA06-4. PCR products were submitted to restriction with the DpnI enzyme (NEB, R0176S) following the manufacturer’s protocol. Both products were purified using GFX™ PCR DNA or Gel Band Purification Kit (GE-healthcare). The NEBuilder HiFi DNA Assembly Cloning Kit (NEB, E5520S) was used to assemble the fragments into pMGAL-Spacer-DR, pMGAL-Spacer-S6-1, pMGAL-Spacer-S6-2, pMGAL-Spacer-CA06-3, pMGAL-Spacer-CA06-4*.* Two µL of the assembly was transformed into *E. coli* NEB 5-alpha (NEB, C2987H). Transformants were screened after DNA minipreps using NucleoSpin Plasmid kit (Macherey-nagel, 740588.50) and enzymatic digestion. Sanger sequencing was performed for final verification.

Plasmid pTi1.0 and 1.1. DNA cassette coding for sgRNA and containing Spiraline promotor (pSpi), two BseRI sites and sgRNA backbone, was constructed. Using PCR Q5 High-Fidelity DNA Polymerase (M0491) kit pSpi and sgRNA sequence were amplified with F35/F36 and F37/F38 primers. Overlap PCR were then achieved between the two fragments and sgRNA cassette was amplified with F35/F38 using Q5® High-Fidelity DNA Polymerase (M0491) kit. pMGAL and sgRNA cassette were digested by EcoRI-HF enzyme (NEB) 2h at 37°C. Digested pMGAL were dephosphorylated by antartic phosphatase kit (NEB M0289S) 2h at 37°C. After purification of the two fragments, ligation was made with T4 DNA ligase (promega) over night at 4°C. Transformation of 2µl of reaction mix was achieved in NEB® 5-alpha Competent *E. coli* (NEB, C2987H). Colonies were screened and plasmid was sequenced and pTI1.0 plasmid was created. Finally, target sequence was added on the different plasmid. Target DNA fragment was prepared by hybridization of two primers: F39/F40, 10µL of each primer (Ci = 100 µM) were heated with 2µL of Adv2 polymerase buffer (TAKARA, 639232) at 95°C, 5min and cooled down slowly (-0,1°C/sec) until room temperature. Plasmids pTI1.0 was digested by BseRI enzyme (NEB R0581S) 2h at 37°C and after purification, targets sequences and plasmids were ligated with T4 DNA ligase (Promega) over night at 4°C. Transformation of 2µl of reaction mix was achieved in NEB® 5-alpha Competent *E. coli* (NEB, C2987H). Colonies were screened and plasmid was sequenced and pTI1.1 plasmid was created.

Maps of plasmids pTi1.0 and pMGAL-Spacer-DR are provided in SI-4.

**PAM library validation.** The 7N degenerated PAM plasmid library was provided by Dr. Gasiunas from CasZyme (Gasiunas et al., 2020). Randomness of the 7 positions of the randomized PAM was validated as described in (Karvelis et al., 2019). Briefly, triplicate of short PCR of 15 cycles were performed on 2.5 ng of pTZ57-7n library using Phusion High-Fidelity DNA Polymerase kit (ThermoFisher, D-530XL) and primers C9/C10. After purification of triplicate pooled product using GeneJET PCR Purification Kit (ThermoFisher, K0701) new step of short PCR of 10 cycles was performed in triplicates using primers C11/C12 and pooled again. PCR products from 3 replicates were sequenced on an Illumina MiSeq V2 (2x150 bp). Datasets treatment was achieved using the tools available at <https://usegalaxy.eu/>. After trimming with Trimmomatic (Galaxy Version 0.36.5, average quality required: Q20) and quality check, the 3 datasets included 1.1 to 1.3 M of paired reads. For each dataset, reads R1 and R2 were merged using FLASH (Galaxy Version 1.2.11.4, Minimum overlap: 50, Maximum overlap: 65, Maximum mismatch density: 0.25) and filtered to perfectly match the expected sequence CGGCGACGTTGGGTCAACTNNNNNNNTGTCCTCTTCCTCTTTAGCGTTTA. Sequence logos were generated with the Sequence Logo software (Galaxy Version 3.5.0) and frequency matrices were calculated for each of the 7 randomized positions. Out of the 16384 possible PAM sequences, 16306 were detected in one or several replicates with a number of reads varying from 1 to 420 (Figure S3A, Table S6A). Base frequency at each of the 7 positions was calculated showing nearly perfect reproducibility among the replicates and observed average frequencies of A, C, G and T ranging from 0.16 to 0.35 over the 7 positions (Figure S3B, Table S6B). The average frequency of each represented PAM was calculated (Figure S3C) and a normalization factor N was calculated as the ratio between the average observed frequency and the theoretical frequency (1/16384) (Table S6A).

***In vitro* cleavage assays.** Codon optimized sequence of MgCas9 strain CA06 under the T7 promotor and T7 terminator was provide by Dr. Gasiunas from CasZyme (Gasiunas et al., 2020). The pET28_MgalS6Cas9 plasmid with codon-optimized sequence of MgCas9 strain S6 was constructed in the lab. A PCR was performed using Advantage HF 2 PCR Kit (Takara, 639123) to produce linear matrice containing sgRNA from pET28-sgRNA plasmid. Cas9 was produced using PURExpress bacterial IVT kit (New England Biolabs). 0.5 µg of MgCas9 plasmid and a 100-fold molar excess of sgRNA linear template was added to the IVT reaction mix. 1 µL of RiboLock RNase Inhibitor (40 U; Thermo Fisher Scientific) were added in the IVT reaction mix. After 4 h of incubation, reaction was stopped by fast cooling on ice. 5 µL of the resulting Cas9-sgRNA ribonucleoprotein (RNP) complex were then combined with 1 μg of plasmid containing target in a 100 µl reaction buffer (10 mM Tris-HCl pH 7.5 at 37°C, 100 mM NaCl, 10 mM MgCl2, 1 mM DTT). After one hour of clivage, DNA were purified using GFX™ PCR DNA or Gel Band Purification Kit (GE-healthcare). Results were observed using Gel electrophoresis.

For the PAM degenerated plasmid library, assays were performed as described in (Karvelis et al., 2019). Using MgCas9 S6 and CA06, 1 µg of 7N Degenerated plasmid library was cleaved. The complete digestion reactions were incubated with 2.5 U of DreamTaq DNA polymerase (5u/µL Thermo Fisher Scientific) and 0.5 μL of 10 mM dATP for an additional 30 min. at 72 °C. DNA were purified with GeneJET PCR Purification Kit. Two oligonucleotides adapters C13/C14 were annealed in primer annealing buffer (10 mM Tris-HCl, 100 mM NaCl, 1 mM EDTA, 1 mM DTT, pH 7.5) during 95°C for 5min and cooling temperature until 25°C. 100 ng of the resulting adapter was ligated to an equal concentration of the purified 3′ dA overhanging cleavage products for 1 h at 22°C in a 25 μL reaction volume in ligation buffer (40 mM Tris–HCl pH 7.8 at 25°C, 10 mM MgCl2, 10 mM DTT, 0.5 mM ATP, 5 % (w/v) PEG 4000, and 0.5 U T4 Ligase; Thermo Fisher Scientific). Using 10 μL of ligation reaction mixtures as a template and primers C13/C16, PCR were performed with Phusion High-Fidelity DNA Polymerase (Thermo Fisher Scientific) for 15 cycles amplification in 100 μL total volume. Purification using GeneJET PCR Purification Kit and a second PCR using primer C11/C15 were performed.

For each MgCas9, 3 independent cleavage assays were performed. After capture of cleaved molecules by adapter ligation and PCR amplification, products were submitted to deep sequencing and datasets were analyzed as previously. Distributions of average PAM frequencies after cleavage by CA06 and S6 MgCas9 were calculated and the top 1000 more frequent PAM sequences (~ top 10%) were selected for further analyses to reduce the impact of background noise generated by rare or non-specific cleavage by the IVT mixture (Figures S4C and S4D). Normalized frequencies were calculated as the ratio between the average observed frequencies after cleavage and the corresponding normalization factor N. PAM frequency matrices and sequence logos were then generated as previously.

As a control of the MgCas9 cleavage specificity, we checked the cleavage frequency of the PAM motif corresponding to the 7 first positions (GTTTTAG) of the Direct Repeat sequence found in the natural CRISPR arrays of *M. gallisepticum* and that is not supposed to be recognized by MgCas9. While the GTTTTAG motif was the 497^th^ most represented motif in the initial PAM library, it was not detected in any of the replicates after cleavage with MgCas9 CA06 or MgCas9 S6. This result suggested the high specificity of the *in vitro* cleavage with IVT mixtures.

**Colony analysis.** PCR screening was performed on all transformants using Advantage HF 2 PCR Kit (Takara, 639123). List of all primers used are detailed in Table S1.

**SI-2. Supplementary results**

**CRISPR locus is part of a putative defense island in *M. gallisepticum* genome.** Detection of additional defence systems in *M. gallisepticum* genomes was performed using PADLOC software (Payne et al., 2021). Three others systems were predicted, including two Restriction-Modification systems (RM) and one Toxin-Antitoxin (TA) system (Table S7). The RM1 that is found in 6 poultry strains close to the CRISPR locus is missing from the genomes of all house finch strains and another type I RM (hereafter named RM2) is found in two poultry strains but not in house-finch strains (Table S7). The TA system *IetA-IetS* has been recently characterized as a bacterial defence systems against phages (Gao et al., 2020) and described as functional in *Mycoplasma mycoides* subsp. *capricolum* and other mycoplasmas (Hill et al., 2021)*.* RM1 and *IetA-IetS* are found in the vicinity of the CRISPR-Cas locus suggesting that the CRISPR-Cas system might be part of a more complex defence island (Figure S6). Two genes encoding another putative S8 serine peptidase (22% coverage, 29% identity with S8 LetAS peptidase) and a hypothetical protein, were also located upstream from the CRISPR locus. S8 serine peptidases are usually found in defence systems against phages (Gao et al., 2020) or sometimes against host immune system (McKenna et al., 2021). This putative S8 peptidase gene was predicted inactivated in 6 poultry strains but functional in all house finch strains except those where the set of *cas* genes had been lost, suggesting both loci were lost during a single deletion event. In addition, a transposase gene of an insertion sequence element (IS256) located immediately downstream of the defence island was conserved in all strains. Concerning RM2, this system is found close to another dynamic genome locus containing genes encoding variable surface proteins (VlhA) (Liu et al., 2000; Pflaum et al., 2015) (Table S7). Therefore, most defence systems against phages in *M. gallisepticum* genomes are cauterized in a highly dynamic putative defence island.

**SI-3. Supplementary discussion**

**Comparison of *in vitro* and *in vivo* findings.** Differences in the repertoires of PAM sequences recognized by the MgCas9 of poultry and house finch isolates were particularly marked in the *in vitro* assay when using a library of plasmids containing all possible combinations of PAM sequences. When comparing these *in vitro* findings with those from an *in vivo* assay based plasmid interference, we confirmed that a lack of recognition of the PAM *in vitro* is associated with a lack of cleavage (and thereby the maintenance) of the plasmid *in vivo*, with the recognition specificity dependent on the PI domain of MgCas9. Indeed, a plasmid harbouring a target sequence followed by the PAM CA06_3 (TTGACTG) was efficiently transformed in *M. gallisepticum* S6 strain, but not in a S6 strain which MgCas9 PI domain had been replaced by the homologous domain originating from CA06 strain MgCas9 (Figure 4). This indicates an efficient cleavage of the plasmid by the engineered MgCas9, in complete accordance with the *in vitro* results.

It is worth noting that some PAM sequences that were only poorly recognized *in vitro* could give rise to sufficient plasmid cleavage to stop colonies from growing and therefore, low numbers of transformants in plasmid interference assays. Such a difference between the *in vitro* and *in vivo* assays can be explained because the highly sensitive *in vitro* assay is designed to quantify cleavage efficiency of PAM sequences after one hour of incubation with the MgCas9/sgRNA complex. In fact, the *in vitro* assay shows quantitative variations in the PAM sequences’ abilities to thermodynamically interact with the MgCas9 PI domain. The *in vivo* assay is based on the cleavage of a low-copy number *oriC* plasmid by the endogenous CRISPR-Cas system of *M. gallisepticum*. Even if not frequent, any cleavage of the plasmid occurring before division of the bacteria will hamper the propagation of the plasmid in the population and stop colony development on selective medium. Beyond the present study, further work *in vivo* would be necessary to determine the actual protection provided by the two MgCas9 against populations of phage harbouring protospacer sequences followed by PAM differentially recognized *in vitro*.

**SI-4. Plasmid maps**

**pTI1.0 :**

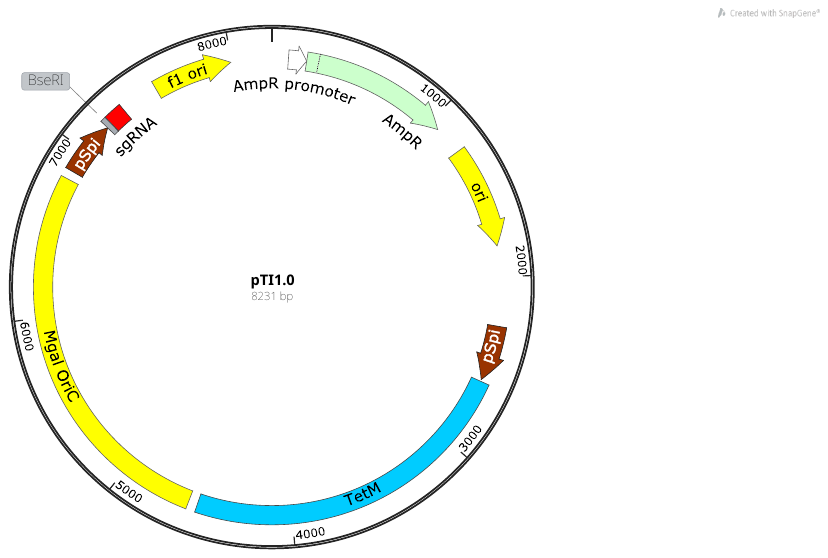

**pMGAL-Spacer-DR:**

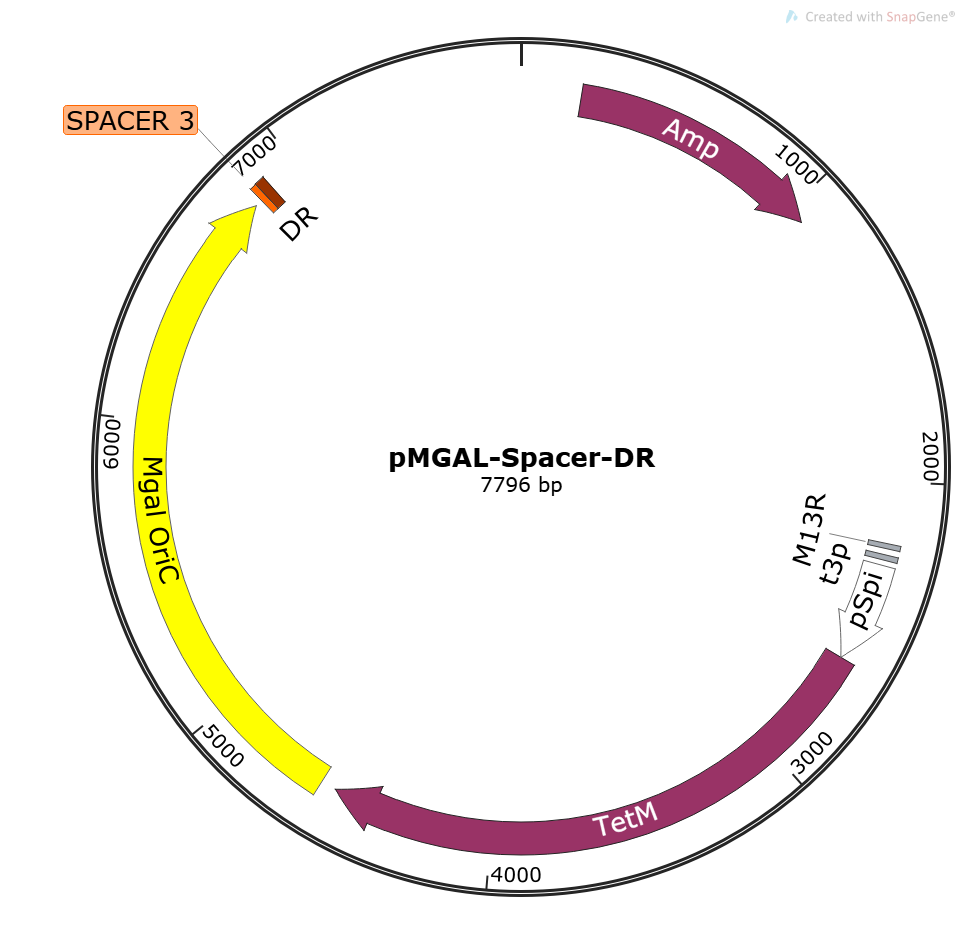

**SI - References**

Arndt, D., Grant, J. R., Marcu, A., Sajed, T., Pon, A., Liang, Y., & Wishart, D. S. (2016). PHASTER: a better, faster version of the PHAST phage search tool. *Nucleic Acids Research*, *44*(W1), W16–W21. https://doi.org/10.1093/nar/gkw387

Chen, I.-M. A., Chu, K., Palaniappan, K., Pillay, M., Ratner, A., Huang, J., Huntemann, M., Varghese, N., White, J. R., Seshadri, R., Smirnova, T., Kirton, E., Jungbluth, S. P., Woyke, T., Eloe-Fadrosh, E. A., Ivanova, N. N., & Kyrpides, N. C. (2019). IMG/M v.5.0: an integrated data management and comparative analysis system for microbial genomes and microbiomes. *Nucleic Acids Research*, *47*(D1), D666–D677. https://doi.org/10.1093/nar/gky901

Darling, A. C. E., Mau, B., Blattner, F. R., & Perna, N. T. (2004). Mauve: Multiple alignment of conserved genomic sequence with rearrangements. *Genome Research*, *14*(7), 1394–1403. https://doi.org/10.1101/gr.2289704

Gao, L., Altae-Tran, H., Böhning, F., Makarova, K. S., Segel, M., Schmid-Burgk, J. L., Koob, J., Wolf, Y. I., Koonin, E. V., & Zhang, F. (2020). Diverse Enzymatic Activities Mediate Antiviral Immunity in Prokaryotes. *Science*. https://doi.org/10.1126/science.aba0372.Diverse

Gasiunas, G., Young, J. K., Karvelis, T., Kazlauskas, D., Urbaitis, T., Jasnauskaite, M., Grusyte, M. M., Paulraj, S., Wang, P., Hou, Z., Dooley, S. K., Cigan, M., Alarcon, C., Chilcoat, N. D., Bigelyte, G., Curcuru, J. L., Mabuchi, M., Sun, Z., Fuchs, R. T., … Siksnys, V. (2020). A catalogue of biochemically diverse CRISPR-Cas9 orthologs. *Nature Communications*. https://doi.org/10.1038/s41467-020-19344-1

Hill, V., Akarsu, H., Barbarroja, R. S., Cippà, V. L., Kuhnert, P., Heller, M., Falquet, L., Heller, M., Stoffel, M. H., Labroussaa, F., & Jores, J. (2021). Minimalistic mycoplasmas harbor different functional toxin-antitoxin systems. *PLOS Genetics*, *17*(10), e1009365. https://doi.org/10.1371/journal.pgen.1009365

Jinek, M., Chylinski, K., Fonfara, I., Hauer, M., Doudna, J. A., & Charpentier, E. (2012). A programmable dual-RNA-guided DNA endonuclease in adaptive bacterial immunity. *Science*, *337*(6096), 816–821. https://doi.org/10.1126/science.1225829

Karvelis, T., Young, J. K., & Siksnys, V. (2019). A pipeline for characterization of novel Cas9 orthologs. In *Methods in Enzymology* (1st ed., Vol. 616). Elsevier Inc. https://doi.org/10.1016/bs.mie.2018.10.021

Kumar, S., Stecher, G., Li, M., Knyaz, C., & Tamura, K. (2018). MEGA X: Molecular evolutionary genetics analysis across computing platforms. *Molecular Biology and Evolution*, *35*(6), 1547–1549. https://doi.org/10.1093/molbev/msy096

Lee, S.-W., Browning, G. F., & Markham, P. F. (2008). Development of a replicable oriC plasmid for *Mycoplasma gallisepticum and Mycoplasma imitans*, and gene disruption through homologous recombination in M. gallisepticum. *Microbiology (Reading, England)*, *154*(Pt 9), 2571–2580. https://doi.org/10.1099/mic.0.2008/019208-0

Li, W., & Godzik, A. (2006). Cd-hit: a fast program for clustering and comparing large sets of protein or nucleotide sequences. *Bioinformatics*, *22*(13), 1658–1659. https://doi.org/10.1093/bioinformatics/btl158

Liu, L., Dybvig, K., Panangala, V. S., Van Santen, V. L., & French, C. T. (2000). GAA trinucleotide repeat region regulates M9/pMGA gene expression in *Mycoplasma gallisepticum*. *Infection and Immunity*, *68*(2), 871–876. https://doi.org/10.1128/IAI.68.2.871-876.2000

McKenna, S., Huse, K. K., Giblin, S., Pearson, M., Majid Al Shibar, M. S., Sriskandan, S., Matthews, S., & Pease, J. E. (2021). The Role of Streptococcal Cell-Envelope Proteases in Bacterial Evasion of the Innate Immune System. *Journal of Innate Immunity*, 1–20. https://doi.org/10.1159/000516956

Payne, L. J., Todeschini, T. C., Wu, Y., Perry, B. J., Ronson, C. W., Fineran, P. C., Nobrega, F. L., & Jackson, S. A. (2021). Identification and classification of antiviral defence systems in bacteria and archaea with PADLOC reveals new system types. *Nucleic Acids Research*, 1–11.

Pflaum, K., Tulman, E. R., Beaudet, J., Liao, X., & Geary, S. J. (2015). Global changes in *Mycoplasma gallisepticum* phase-variable lipoprotein gene vlhA expression during in vivo infection of the natural chicken host. *Infection and Immunity*, *84*(1), 351–355. https://doi.org/10.1128/IAI.01092-15
